# Mechanistic model for epigenetic maintenance by methyl-CpG-binding domain proteins

**DOI:** 10.1101/2024.09.22.614380

**Authors:** Liuhan Dai, Alexander Johnson-Buck, Nils G. Walter

## Abstract

DNA methylation is a fundamental element of epigenetic regulation that is governed by the MBD protein superfamily, a group of “readers” that share a highly conserved methyl-CpG-binding domain (MBD) and mediate chromatin remodeler recruitment, transcription regulation, and coordination of DNA and histone modification. Previous work has characterized the binding affinity and sequence selectivity of MBD-containing proteins toward palindromes of 5-methylcytosine (5mC) containing 5mCpG dinucleotides, often referred to as single symmetrically methylated CpG sites. However, little is known about how MBD binding is influenced by the prototypical local clustering of methylated CpG sites and the presence of DNA structural motifs encountered, e.g., during DNA replication and transcription. Here, we use Single-Molecule Kinetics through Equilibrium Poisson Sampling (SiMKEPS) to measure precise binding and dissociation rate constants of the MBD of human protein MBD1 to DNAs with varying patterns of multiple methylated CpG sites and diverse structural motifs. MBD binding is promoted by two major properties of its DNA substrates: 1) tandem (consecutive) symmetrically methylated CpG sites in double-stranded DNA and secondary structures in single-stranded DNA; and 2) DNA forks. Based on our findings, we propose a mechanistic model for how MBD proteins contribute to epigenetic boundary maintenance between transcriptionally silenced and active genome regions.

## Introduction

In eukaryotes, DNA methylation refers to the addition of a methyl group to a nucleobase—typically to the 5-position of cytosine to form 5-methylcytosine, 5mC—in double-stranded DNA (dsDNA). In mammals, DNA methylation occurs almost exclusively at CpG dinucleotides^1^. Over the past three decades, promoter hypermethylation at CpG islands has been linked to heritable transcriptional repression, and such DNA methylation-mediated gene silencing has been found to play a crucial role in many biological processes such as mammalian development, X chromosome inactivation, genomic imprinting, and genome stability^2–7^.

In mammals, DNA methylation remodels chromatin structure and alters locus-specific gene expression states by recruiting a class of methylation reader proteins, the MBD (methyl-CpG-binding domain) superfamily^8–10^. All MBD proteins share a conserved MBD domain, consisting of approximately 70-85 amino acids, that recognizes 5mCpG motifs^8^. A single MBD domain alone can bind a single symmetrically methylated CpG dinucleotide surrounded by over 12 bp dsDNA^11,12^. Previous studies have focused on characterizing the sequence selectivity and affinity of various MBD domains in the context of binding to single palindromes of 5mCpG dinucleotides^13–16^. However, little is known about how MBD binding to 5mCpGs depends on the common local clustering of methyl modifications characteristic of epigenetic gene silencing or the DNA structural motifs encountered during central cellular processes such as DNA replication and transcription. Filling this knowledge gap is critical for understanding and manipulating key processes governed by MBD proteins in epigenetics, including the recruitment of chromatin modifying complexes, regulation of gene silencing, and prevention of epigenetic mark spread.

Here, we develop Single-Molecule Kinetics through Equilibrium Poisson Sampling (SiMKEPS) into a tool that measures precise binding and dissociation rate constants of the prototypical MBD of human protein MBD1 to DNA. By systematically varying DNA substrates to contain different methylation patterns and structural motifs mimicking replication and transcription intermediates, we discovered two novel features that influence MBD binding kinetics and affinity. First, the presence of tandem symmetrically methylated CpG sites increases the affinity of MBD binding to both dsDNA and secondary structured single-stranded DNA (ssDNA). Second, the presence of the bifurcation in a hemimethylated dsDNA fork allows for stable MBD binding even in the absence of symmetrical CpG methylation. Our results lead to a mechanistic model that offers a comprehensive understanding of how the MBD superfamily proteins recognize and take action on clustered epigenetic marks.

## Results

### Halo-tagged MBD preferentially binds a symmetrically 5-methylated BCAT promoter

The methyl-CpG-binding domain (MBD, aa 1-77) of human MBD1 was fused through a GGGSG linker with a C-terminal HaloTag for site-specific labeling. As expected, the resulting MBD-Halo design was predicted by AlphaFold2^17^ to fold into independent MBD and HaloTag domains connected by the flexible linker (**Fig. 1a**), reducing the likelihood that the HaloTag will interfere with the folding or function of the MBD domain. The methyl-CpG binding activity of purified MBD-Halo was validated by an electrophoretic mobility shift assay (EMSA) at low-ion strength (to stabilize all protein:DNA interactions), visualized by fluorescence staining (**Fig. 1b**). An excess of MBD-Halo was incubated with three exemplary DNA substrates: unmethylated dsDNA (UM), hemimethylated dsDNA (HM), and symmetrically methylated dsDNA (SM). All three DNAs share the same sequence, a 55-bp fragment of the branched-chain amino acid transaminase 1 (BCAT1) promoter of which up to 7 pairs of CpG dinucleotides may be methylated^18,19^. We chose the BCAT promoter since its hypermethylation is a widespread hallmark of cancer, leading to the downregulation of BCAT1 expression, thereby affecting cancer cell metabolism and the tumor microenvironment. A concentration-dependent band shift occurred when MBD-Halo was incubated with each of the three BCAT1 DNA substrates (**Fig. 1b**), indicating that MBD-Halo can bind all substrates when present at sufficiently high concentration. However, while a 10-fold excess of MBD-Halo was sufficient to yield strong binding of SM as evidenced by a prominent shifted band corresponding to the protein-DNA complex, no evidence of binding to HM or UM was evident at this concentration of MBD-Halo. This observation is consistent with expectations that MBD-Halo binds much more strongly to symmetrically methylated dsDNA than to either hemimethylated or unmethylated dsDNA. Furthermore, as the ratio of MBD-Halo to SM was increased from 5-fold to 50-fold, we observed three differentially shifted bands, likely corresponding to complexes with three different protein:DNA stoichiometries: 1:1, 2:1 and 3:1. This observation is consistent with a prior report suggesting a binding footprint of ∼12 bp for MBD^11^, which would only permit binding of up to three MBD-Halos within the ∼39 bp spanned by the methylated CpGs in the 55 bp BCAT1 sequence used here. Despite MBD’s lower affinity for it, the HM substrate also showed evidence of binding to multiple copies of MBD at the highest concentration (50x) of MBD-Halo, as evidenced by at least 2 shifted bands corresponding to protein:DNA ratios of 1:1 and 2:1.

**Fig. 1.**
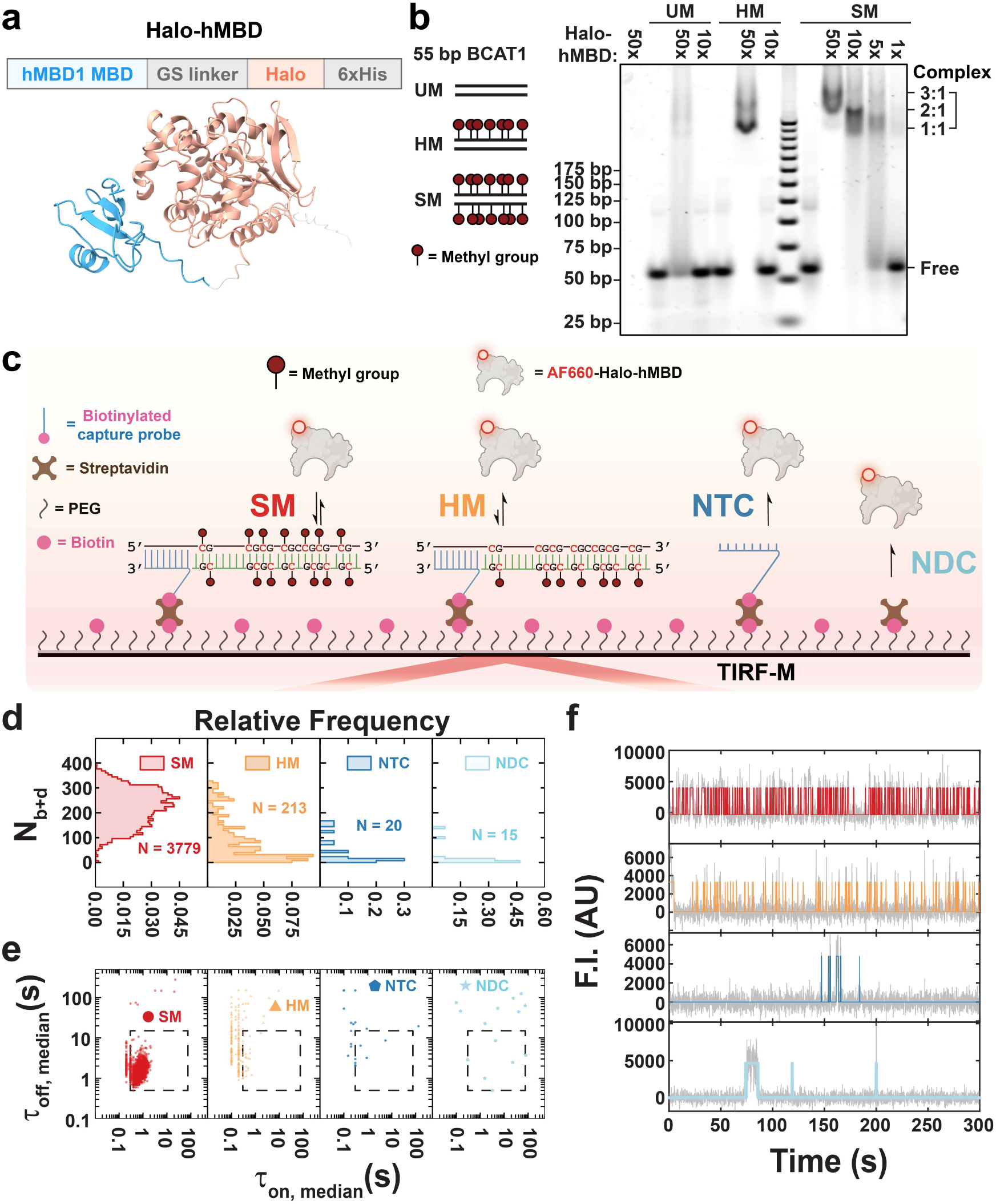
MBD-Halo shows methyl-CpG-binding activity in both ensemble and single-molecule measurements. **a** Predicted structures of MBD-Halo by AlphaFold2. See **Supplementary Table** for input parameterization. **b** EMSA of MBD-Halo binding to three types of 55 bp BCAT1 promoter substrates with different methylation states: SM (symmetrically methylated DNA), HM (hemimethylated DNA), and UM (unmethylated DNA). MBD-Halo was mixed with DNAs at different molar ratios. **c** Schematic of single-molecule fluorescence microscopy for studying MBD binding to surface-tethered methylated DNA. NTC, non-target control; NDC, non-DNA control. **d,e,f** *N_b+d_* distributions (**d**), median dwell time distributions (**e**), and representative intensity-time traces (**f**) under four different conditions: SM, HM, NTC and NDC. N, number of detected molecules of one field of view (FOV). Gray lines and colored lines in panel f represent raw and idealized traces by hidden Markov modeling respectively. F.I., fluorescence intensity; AU, arbitrary unit.

Except at the highest concentrations and stoichiometries of MBD-Halo combined with HM or SM, a tailing effect was observed below the shifted bands, suggesting gradual dissociation of the complex once it entered the gel (**Fig. 1b**). We hypothesized that MBD-Halo’s concentration (0.5 and 1 µM for 5x and 10x MBD-Halo, respectively) fell around its K_D_ for SM while migrating in the gel, leading to gradual dissociation of protein:DNA complex. This is consistent with the relatively high K_D_ values (0.6 ∼ 0.9 µM) for the interaction of MBD with symmetrically methylated DNA measured by Liu *et al*^16^. We further verified methyl-CpG binding activity of Alexa Fluor 660-labeled MBD-Halo (MBD-Halo-AF660) using EMSA under the same buffer conditions (**Supplementary Fig. 1**). Unsurprisingly, since the highest concentration of MBD-Halo-AF660 available was 2x (200 nM)—well below the expected K_D_—we did not observe a distinct band corresponding to the DNA-protein complex (**Supplementary Fig. 1**). However, consistent with our results for MBD-Halo in **Fig. 1B**, we observed a tailing band shift at 1.4x and 2x MBD-Halo-AF660 when incubated with SM but not for UM or HM (**Supplementary Fig. 1**), again suggesting a higher affinity of MBD-Halo-AF660 for symmetrically methylated dsDNA than hemimethylated or unmethylated dsDNA.

Next, we used single-molecule fluorescence microscopy to confirm methylation-specific interactions of MBD-Halo-AF660 (**Fig. 1c**) under the same buffer conditions as in the EMSA. Each target (fully methylated or unmethylated 55-nt ssDNA sense-strand of BCAT1) was introduced at a concentration of 10 pM and immobilized through hybridization with a 5’-biotinylated antisense capture probe with overhang (CPO) that was anchored on a biotin-PEG (polyethylene glycol)-passivated surface *via* a bridging streptavidin and extended ssDNA linker (**Fig. 1c**). A fully methylated antisense strand was hybridized with the remainder of the BCAT1 sense strand to generate either a symmetrically methylated (SM) or hemimethylated (HM) design in separate experiments (**Fig 1c**). We observed MBD-Halo-AF660 binding to each of these dsDNAs under low ionic strength conditions (similar to those in the EMSA) by total internal reflection fluorescence microscopy (TIRF-M, see Methods). In the resulting videos, we identified diffraction-limited spots that showed significantly higher fluorescence fluctuations than their surrounding pixels— corresponding to locations of repeated binding and, hence, likely representing regions containing a single immobilized target molecule—and then calculated the fluorescence intensity over time for each such spot. To examine the non-specific binding of MBD-Halo-AF660 to surface matrix, two negative controls—non-target control (NTC, with CPO but no target) and non-DNA control (NDC, with neither CPO nor target)— were tested separately and showed only a minimal number of spots (**Fig 1c**), allowing us to utilize all surface spots in the experiments with DNA present. Having confirmed MBD-Halo-AF660’s functional activity, we henceforth refer to it simply as “MDB”.

Three key kinetic parameters were extracted from these time traces: *N_b+d_* (average number of binding and dissociation events to a target molecule within the observation window; **Fig. 1d**), *τ_on_* (weighted-average time constant for leaving the MBD-bound state, calculated by exponential fitting of the cumulative distribution of “fluorescence on” times and thus an ensemble-averaged property) and *τ_off_* (similarly the weighted-average time constant for leaving the MBD-unbound state, calculated by exponential fitting of the cumulative distribution of “fluorescence off” times). In addition, *τ_on,median_* and *τ_off,median_* (the median dwell times of individual traces in the on and off states, respectively) were used to represent individual traces (**Fig. 1e**). Consistent with the tailing of bands observed on EMSA (**Fig. 1b**), the binding of MBD to both SM and HM targets was highly reversible, as evident from the repeated patterns of binding and dissociation shown in the time traces (**Fig. 1f**). This observation allowed us to measure the protein binding and dissociation time constants for these two substrates at high precision via Single-Molecule Kinetics through Equilibrium Poisson Sampling (SiMKEPS). Notably, we observed >10-fold more traces (i.e., spots in TIRF-M) for the SM substrate than the HM substrate, defining the N molecules with observable MBD binding and dissociation events (dashed box in **Fig. 1d**), and >100-fold more traces in HM than in either of the two negative control conditions NTC and NDC (**Fig. 1d,e**), with significant differences also evident in representative trajectory patterns (**Fig. 1f**). These observations are consistent with the expectation that MBD will bind most strongly to the symmetrically methylated SM target. Note that we expect similar numbers of SM and HM molecules to be immobilized because they were introduced at the same concentration and share the same DNA sequence; the difference in the number of bona fide traces, N, therefore reflects consistently shorter MBD binding events to HM than to SM, often not passing the threshold for observation at our 100 ms time resolution. (Further evidence is that we observed an accumulation of τ_on,median_ values close to this time resolution for HM, suggesting that shorter events will be missed; **Supplementary Fig. 2a**.) Consistent with these differential binding kinetics, the *N_b+d_* distribution of SM molecules was shifted up (**Fig. 1d**), with an average of N_b+d_ = 220 events per trace, compared to 94 for HM (**Fig. 1e**), with significantly higher *τ_on,median_* and lower *τ_off,median_* values (**Supplementary Fig. 2a** and **Supplementary Fig. 2b**), as expected for more thermodynamically stable MBD binding to the SM than the HM target. Accordingly, *τ_on_* and *τ_off_* for SM differed significantly with values of 0.9 s and 1.6 s, respectively, versus HM’s 0.3 s and 5.2 s, respectively, suggesting that the bound time is extended and the unbound time shortened for SM, both contributing to the higher affinity of MBD for SM than HM (**Supplementary Fig. 2c** and **Supplementary Fig. 2d**).

Taken together, our ensemble EMSA and single-molecule measurements provide evidence of more stable, yet still transient, binding of MBD to symmetrically methylated dsDNA compared to hemimethylated and especially unmethylated targets. The relative affinity of MBD to variously modified DNA molecules is readily assessed by measuring both its binding and dissociation time constants and the total number *N* of DNA molecules that present as spots with significant signal in TIRF-M.

### DNA fork motif stabilizes MBD binding to hemimethylated DNA

In the cell, epigenetic marks dictate the accessibility of the dsDNA genome for transcription, which leads to transient, partial unfolding into ssDNA segments, as do replication and repair processes. We therefore sought to examine how the resulting Y-shaped DNA fork structures influence MBD binding to methylated DNA targets. To this end, we hybridized our positive-sense BCAT1 strand containing 7 methylated sites (M7) with both the CPO and either of two unmethylated antisense strands (**Fig. 2a**, **Supplementary Fig. 3a**): A39, which forms 39 bp with the target DNA in a blunt-ended duplex, or A17O, forming a 17-bp helix with BCAT1 and a 10-nt overhang at its 5’-end, leaving a 22-nt overhang at the 3’-end of the M7 target, forming a bifurcating DNA fork motif. The blunt-ended hemimethylated M7-A39-CPO (the M7 target hybridized to A39 and captured by CPO) showed an *N_b+d_* of 74, a lifetime of 0.4 s for bound state and 6.8 s for unbound state (**Fig. 2a**, **Supplementary Fig. 3a** and **Supplementary Fig. 4a**). These values are quite similar to those of the also hemimethylated HM substrate, which carries the 5mC modifications on the antisense instead of the sense BCAT1 strand (**Fig. 1c-f**). Also similarly to HM, a large fraction of traces for M7-A39-CPO show a *τ_on,median_* close to 0.1 s (**Supplementary Fig. 4a**), suggesting that a significant fraction of binding events exhibit dwell times shorter than our 100 ms time resolution.

**Fig. 2.**
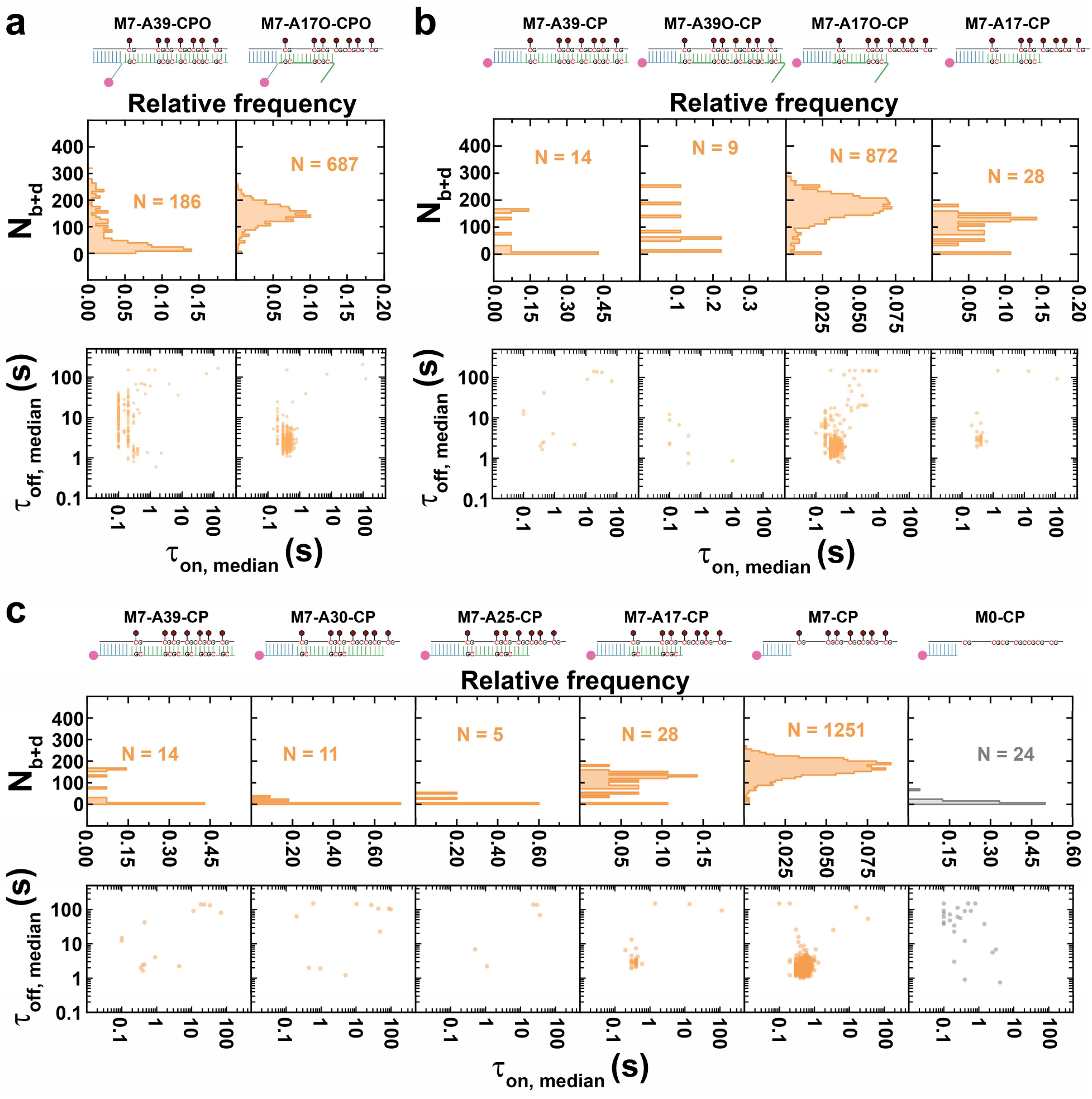
Bifurcation motif stabilizes MBD binding to hemimethylated DNA. **a** Designs of M7-A39-CPO and M7-A17O-CPO as well as their *N*_b+d_distributions and dwell time distributions. **b** Designs of M7-A39-CP, M7-A39O-CP, M7-A17O-CP and M7-A17-CP as well as their *N*_b+d_ distributions and dwell time distributions. **c** Designs of M7-A39-CP, M7-A30-CP, M7-A25-CP, M7-A17-CP, M7-CP and M0-CP as well as their *N*_b+d_ distributions and dwell time distributions. N, number of detected molecules of one FOV.

In striking contrast, the DNA fork-containing design M7-A17O-CPO behaved more similarly to the SM target, with many more detectable molecules, a higher *N_b+d_* (average of 144), as well as longer *τ_on,median_* and shorter *τ_off,median_* than either HM or M7-A39-CPO (**Fig. 2a**, **Supplementary Fig. 3a**, **Supplementary Fig. 4a** and **Supplementary Fig. 4b**). Similarly, *τ_on_* was measured as 0.5 s, and *τ_off_* as 3.2 s, indicating a higher affinity of MBD for this bifurcating complex than the blunt-ended M7-A39-CPO (**Fig S4c** and **Fig S4d**, see **Fig S5** and **Fig S6** for methylation dependency). To test the hypothesis that the DNA fork formed by A17O stabilizes MBD binding to hemimethylated DNA, we designed a series of related BCAT1 DNA targets where the CPO was replaced by a 3’-biotinylated capture probe with no overhang (CP). In addition an overhang of the same sequence as the overhang of A17O was added to the A39 probe, yielding A39O, or the overhang was removed from A17O, yielding A17. Notably, neither M7-A39-CP nor M7-A39O-CP showed detectable MBD binding (**Fig. 2b** and **Fig S3b**), suggesting that the presence of no or only a single 5’-terminal overhang is not sufficient to enhance the affinity of MBD to the level seen with the DNA fork M7-A17V-CPO or even that of M7-A39-CPO, with an internal overhang (compare **Fig. 2b** with **Fig. 2a**). Conversely, these observations suggest that an internal overhang alone can modestly enhance MBD binding to a hemimethylated target (**Supplementary Fig. 5**), whereas a terminal overhang as present in M7-A39O-CP does not (**Supplementary Fig. 2b**).

To further test this hypothesis, we next tested M7-A17O-CP, which uses CP instead of CPO and thus removes the internal overhang of the capture probe relative to M7-A17O-CPO (**Fig. 2b** and **Supplementary Fig. 3b**). In contrast to M7-A39O-CP, M7-A17O-CP exhibited strong MBD binding, again underlining the importance of the DNA fork motif, and signaling the importance of both methylated and juxtaposed unmethylated ssDNA overhangs for stabilization of MBD binding. Like M7-A17O-CPO (**Fig. 2a**), and in contrast to M7-A39O-CP (**Fig. 2b**), M7-A17O-CP exhibits a high *N_b+d_* (average of 166), a relatively long *τ_on_* of 0.6 s, and a relatively short *τ_off_* of 2.4 s (**Supplementary Fig. 4c** and **Supplementary Fig. 4d**), further supporting this conclusion; the internal overhang seems of little importance. To test whether a methylated ssDNA overhang alone can stabilize MBD binding, we tested M7-A17-CP, which lacks the juxtaposed unmethylated overhang; this design shows minimal interaction with MBD (**Fig. 2b** and **Supplementary Fig. 3b**).

Taken together, these results suggest that two adjacent terminal ssDNA overhangs—one methylated, the other unmethylated—jointly act to cooperatively stabilize MBD binding to hemimethylated DNA, and that MBD binding is weak or absent for single terminal (methylated or unmethylated) overhangs.

### An extended ssDNA binds MBD in a methylation and secondary-structure dependent manner

To examine the potential of methylated ssDNA to bind MBD directly, we tested a series of designs with varying lengths of methylated BCAT1 M7 target that lacks overhangs on the antisense strand: M7-A30-CP, M7-A25-CP and M7-CP in addition to M7-A39-CP and M7-A17-CP (**Fig. 2c** and **Supplementary Fig. 3c**). As expected, M7-A39-CP, M7-A30-CP, M7-A25-CP and M7-A17-CP—containing ssDNA overhangs of 0-22 nt with 0-4 methylated CpG sites—showed little to no detectable binding of MBD (**Fig. 2c** and **Supplementary Fig. 3c**), consistent with their lack of an overhang on the juxtaposed antisense strand to produce a DNA fork motif. Strikingly, upon further exposing the methylated target strand in M7-CP (39 nt ssDNA overhang with 7 methylation sites), stable MBD binding was observed with an *N_b+d_* of 167, a relatively long *τ_on_* of 0.7 s, and a relatively short *τ_off_* of 2.5 s (**Fig. 2c**, **Supplementary Fig. 3c**, **Supplementary Fig. 4c** and **Supplementary Fig. 4d**), resembling the values of M7-A17O-CP (**Fig. 2b**). This interaction is methylation-dependent, as it is completely absent from the corresponding unmethylated design M0-CP (**Fig. 2c**). These results suggest that, unexpectedly, a fully exposed methylated BCAT1 ssDNA alone can be stably bound by MBD.

Since methylation is necessary for MBD binding to ssDNA, we asked whether the positioning and number of methylation sites affect MBD binding. To this end, we tested a series of target ssDNAs with different patterns of 5mC modifications, including: M7-CP, M4a-CP, M3b-CP, M3a-CP, M2c-CP, M2b-CP, M2a-CP, M1b-CP and M1a-CP, where the number after ‘M’ represents the number of 5-methyl-CpGs (**Fig. 3a**, **Supplementary Fig. 7a**). Notably, stable MBD binding was only observed to targets with at least 4 methylation sites (**Fig. 3a-c** and **Supplementary Fig. 7a**), suggesting that four is the minimum number of methylation sites to recruit MBD to ssDNA. In contrast, the results in **Fig. 2** indicate that the presence of even seven methylation sites in one strand of a double-stranded target without a DNA fork do not significantly stabilize MBD binding. Combining these observations, we hypothesized that a secondary structure may be responsible for the enhanced MBD binding observed to an extended, multiply methylated ssDNA. The G/C-rich BCAT1 sense strand is predicted to adopt several secondary structures (**Supplementary Fig. 8**), with the most abundant conformer comprising a stem-loop including a tandem symmetrically methylated, 5-bp duplex (**Fig. 3d, Fig S8a**). To test our hypothesis, we designed a new strand, M4b, with also four methylation sites, one of which is moved from the stem to the loop (**Fig. 3e**). Strikingly, this seemingly small change almost completely eliminated MBD binding (**Fig. 3f** and **Supplementary Fig. 7b**), suggesting that the placement of the fourth 5mC on the stem is critical. We further disrupted the stem-loop by hybridizing M4a with increasingly longer antisense strands A39, A30, A25, or A17 (**Fig. 3f** and **Supplementary Fig. 7b).** As expected, these designs generally showed no significant MBD binding (**Fig. 3f**), with the exception of a small population of M4a-A17-CP molecules, for which it is possible that the dsDNA ends fray due to competition with the proposed secondary structure within the sense strand.

**Fig. 3.**
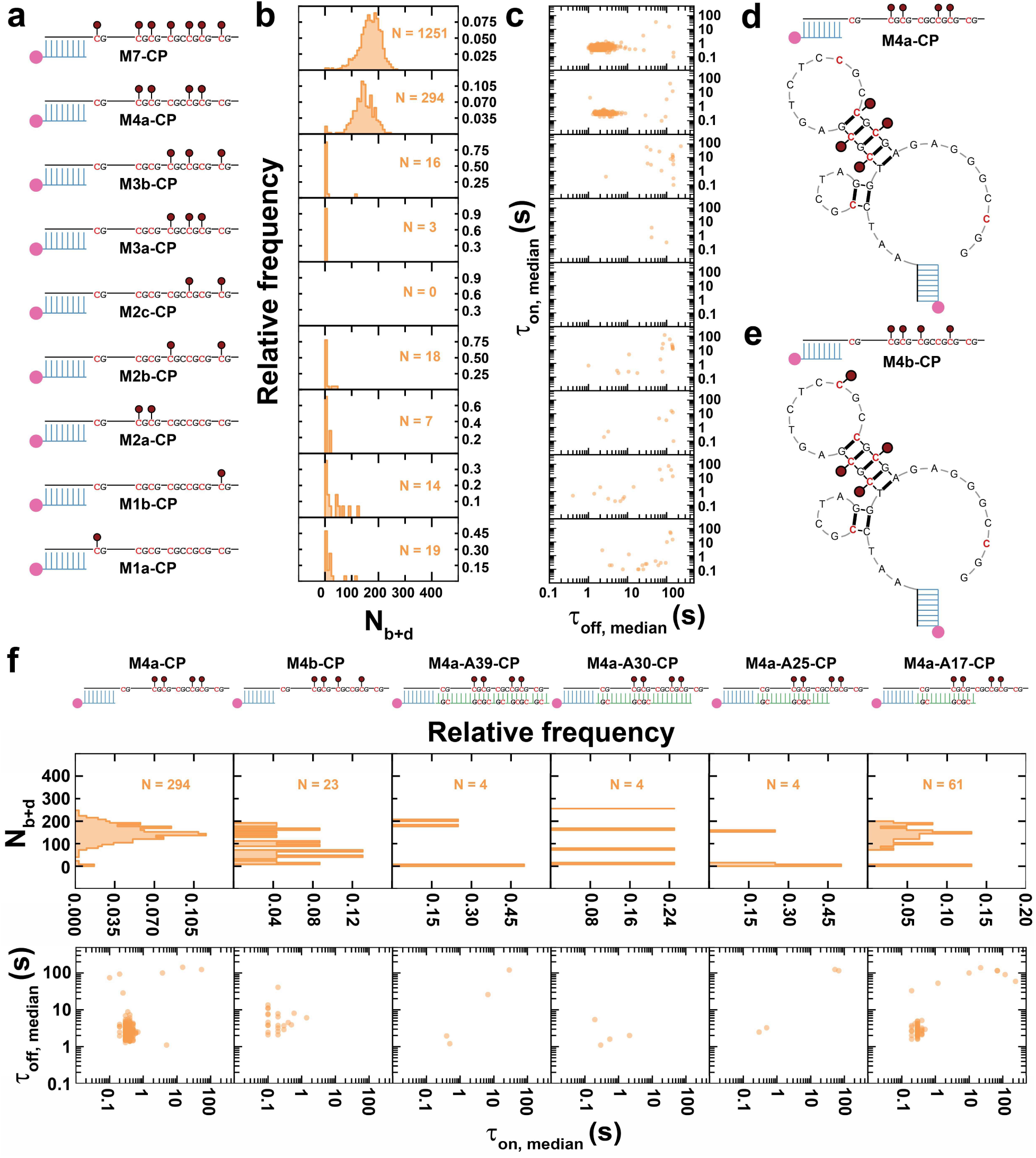
Tandem symmetrically methylated stem loop stabilizes MBD binding to methylated ssDNA. **a-c** Designs of M7-CP, M4a-CP, M3b-CP, M3a-CP, M2c-CP, M2b-CP, M2a-CP, M1b-CP and M1a-CP, as well as their *N*_b+d_ distributions and dwell time distributions. **d-e** Predicted secondary structures of M4a-CP and M4b-CP. **f** Designs of M4a-CP, M4b-CP, M4a-A39-CP, M4a-A30-CP, M4a-A25-CP and M4a-A17-CP, as well as their *N*_b+d_ distributions and dwell time distributions. N, number of detected molecules of one FOV.

These results suggest that a stem-loop secondary structure containing at least two symmetrically methylated tandem CpG sites is a key motif enabling MBD binding to ssDNA. Notably, in this context MBD does not require a contiguous ≥12 helical footprint for stable binding^11^ (**Fig. 3d**), perhaps due to electrostatic attraction by neighboring secondary structures. While also lacking a designed DNA fork motif, the predicted secondary structure arguably resembles a fork with a dsDNA segment flanked by two juxtaposed overhangs (**Fig. 3d**).

While a minimum of two symmetrically methylated tandem CpG sites are sufficient for MBD binding in the context of a secondary structured ssDNA, increasing to a total of seven 5mCs in the BCAT1 sense strand still increases the frequency and duration of binding events further. This is evident from M4a-CP showing a lower *N_b+d_* of 143, a smaller *τ_on,median_*, a larger *τ_off,median_*, and ∼4-fold fewer detected molecules N compared to M7-CP (**Fig. 3a**, **Supplementary Fig. 7a**, **Supplementary Fig. 9a** and **Supplementary Fig. 9b**); M4a-CP also exhibited a ∼25% shorter *τ_on_* (0.5 s) and a ∼36% longer *τ_off_* (3.3 s) than M7-CP (**Supplementary Fig. 9c** and Supplementary Fig. 9d). As previously noted, M4a-CP can adopt multiple secondary structures so that it is possible that only a (majority) fraction of immobilized molecules fold into the most stable conformer that contains the tandem symmetrically methylated stem loop (**Fig. 3d**, **Fig S8a**) while the rest either stay in single-stranded form or adopt other conformations; as a result, only a minority of surface-immobilized targets may exhibit detectable binding of MBD. While this is also true of M7-CP, its larger number of methylation sites combined with the presence in several of the alternative secondary structures of ‘DNA fork’ motifs, already shown to stabilize MBD binding, may render a larger fraction of M7-CP molecules capable of detectably binding MBD. Furthermore, it has long been known that 5-methyl cytosine increases the melting temperature of duplex DNA through increased stability of base stacking^20,21^; the additional methyl groups of M7-CP may thus favorably stabilize symmetrically methylated stem loops relative to single-stranded or other conformations, and result in a higher fraction of conformations with sites capable of MBD binding. Furthermore, the slightly longer dwell times in the MBD-bound state of M7-CP may arise from additional transient interactions with the added methyl groups, which may not only transiently form symmetrically methylated CpG stems but also increase the hydrophobic contact area.^22^

### Interplay between ssDNA overhangs and secondary structures

To better understand the relative importance and possible interplay of overhangs and secondary structures in the BCAT1 sense strand for MBD binding, we next tested a series of designs with an overhang-containing capture probe hybridized to methylated targets with extended ssDNA segments: M7-CPO, M4a-CPO, M3b-CPO, M3a-CPO, M2c-CPO, M2b-CPO, M2a-CPO, M1b-CPO and M1a-CPO, M0-CPO and CPO (**Supplementary Fig. 10**). These designs produce a bifurcation motif while also being able to form secondary structures that, based on their predicted *ΔG* (**Supplementary Fig. 8**), are expected to be favored. Just as with the designs lacking a CP overhang (**Fig. 3a**), only when the number of methylation sites reaches 4 or higher is MBD binding observed (**Supplementary Fig. 10**). Compared to M7-CP, M7-CPO with an additional central overhang exhibits a similar number *N* of detectable molecules with an *N_b+d_* of 180 and a similar *τ_off,median_*, but shows a slight yet significant (p < 0.01) decrease in the individual molecule *τ_on,median_* (**Supplementary Fig. 11a** and **Supplementary Fig. 11b**). Consistent with this observation, M7-CPO showed a shorter *τ_on_* of 0.4 s (M7-CP: 0.7 s) and an unchanged *τ_off_* of 2.5 s (**Supplementary Fig. 11c** and **Supplementary Fig. 11d**). These results suggest that, in the presence of the BCAT1 sense strand secondary structure, the overhang no longer cooperatively stabilizes MBD binding as it does in the DNA fork motifs (**Fig. 2**, M7-A17O-CP and M7-A17-CP). Similarly indicating a weaker MBD binding affinity, M4a-CPO showed a lower number *N* of detectable molecules with an *N_b+d_* of 106, no significant difference *τ_on,median_* and a slight but significant (p < 0.01) increase in *τ_off,median_* compared to M4a-CP (**Supplementary Fig. 11a** and **Supplementary Fig. 11b**); accordingly, M4a-CPO showed an unchanged *τ_on_* of 0.5 s and a longer *τ_off_* of 4.5 s (M4a-CPO: 3.3 s; **Supplementary Fig. 11c** and **Supplementary Fig. 11d**). As for M7-CPO, these results show no additional stabilization of MBD binding upon introduction of an overhang in M4a-CPO, in contrast to M4a-A17O-CP where the overhang serves as part of a DNA fork with no predicted secondary structure and stabilizes MBD binding (**Supplementary Fig. 12**, **Supplementary Fig. 13** and **Supplementary Fig. 14**).

Taken together, these results suggest that secondary structure competes with stabilization of MBD binding to a bifurcated DNA fork motif, and that these two modes of stabilization do not act cooperatively.

### Symmetrically methylated tandem CpG sites stabilize MBD binding to dsDNA

Based on the above observations that MBD binding is highly sensitive to the number and positioning of CpG methylation sites in hemimethylated ssDNA, we expected to observe a similar modulation of MBD binding to symmetrically methylated dsDNA. To investigate this hypothesis, we tested a series of wholly double-stranded designs with a fully antisense strand (A39m): M7-A39m-CP, M4a-A39m-CP, M3b-A39m-CP, M3a-A39m-CP, M2c-A39m-CP, M2b-A39m-CP, M2a-A39m-CP, M1b-A39m-CP, and M1a-A39m-CP **(Fig. 4, Supplementary Fig. 15** and **Supplementary Fig. 16).**

**Fig. 4.**
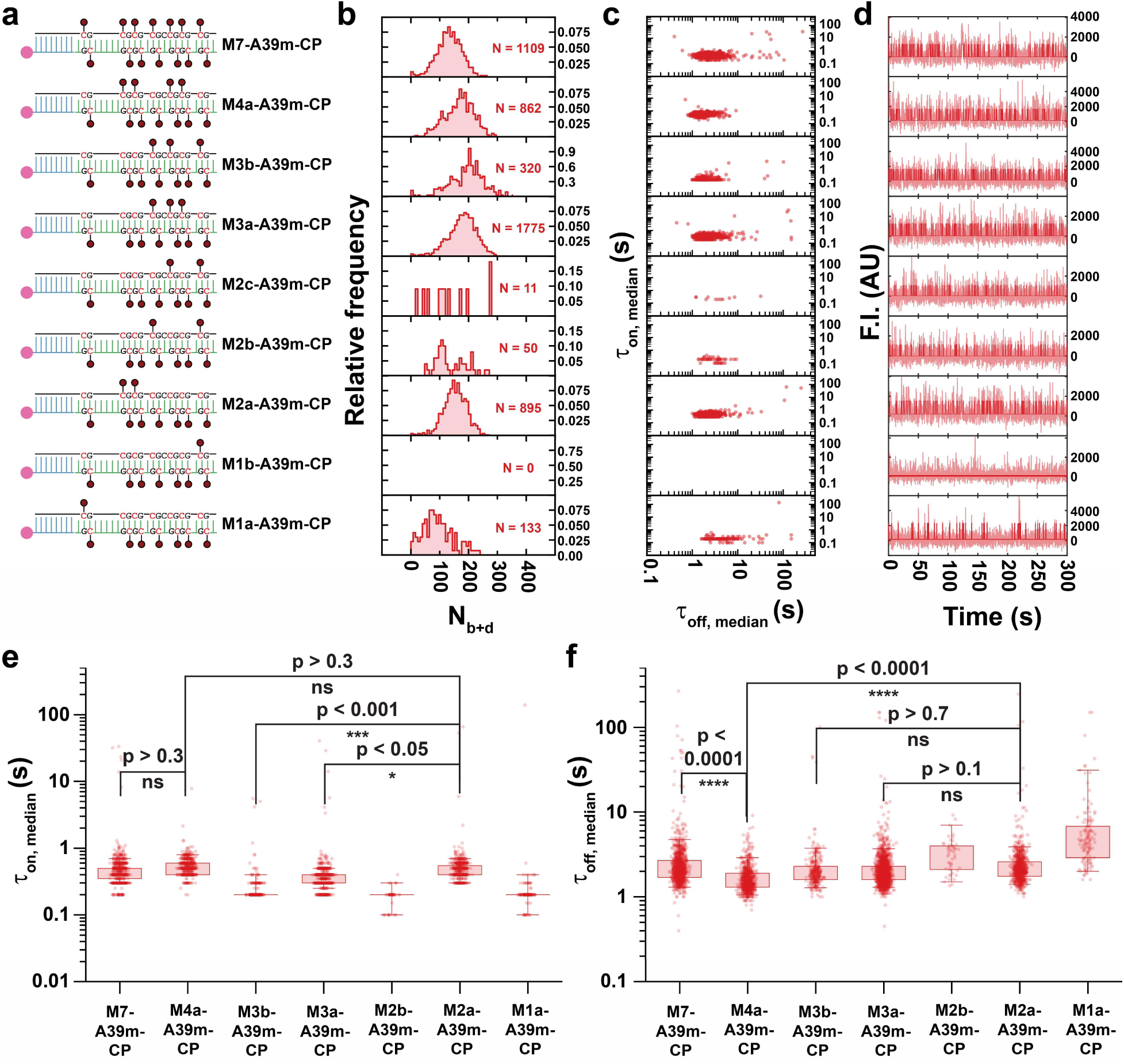
Tandem symmetrical methylation boosts methyl-binding activity of MBD in methylated duplex DNA. **a-d** Designs of M7-A39m-CP, M4a-A39m-CP, M3b-A39m-CP, M3a-A39m-CP, M2c-A39m-CP, M2b-A39m-CP, M2a-A39m-CP, M1b-A39m-CP, and M1a-A39m-CP, as well as their *N*_b+d_distributions, dwell time distributions and representative intensity-time traces. In panel d, semi-transparent lines in the background are raw traces and solid lines are idealized traces by hidden Markov modeling. F.I., fluorescence intensity; AU, arbitrary unit. **e,f** Boxplot comparison of τ_on,median_ and τ_off,median_ distributions respectively, across designs showing significant detected population. Boxes are drawn from Q1 to Q3 with whiskers from 5% percentile to 95% percentile. P-values smaller than 0.05 are assessed using a single-tailed unpaired t-test and P-values higher than 0.05 are assessed using a two-tailed unpaired t-test.

In two cases the BCAT1 sense strand carried only one methylation, yielding a mostly hemimethylated duplex with a single symmetrically methylated CpG site: M1b-A39m-CP, whose symmetrically methylated site is near the end of the duplex; and M1a-A39m-CP, where the site is internal. Notably, while M1b-A39m-CP yielded no detectable MBD binding, M1a-A39m-CP yielded a significant population of ∼133 molecules exhibiting repeated MBD binding, with an *N_b+d_* of 94 (**Fig. 4**). These results suggest that a terminal methyl-CpG (3 bp away from the 3’-end of the sense strand in M1b-A39m-CP) does not provide as stable a binding site for MBD as the internal methyl-CpG of M1a-A39m-CP, possibly due to dsDNA end fraying under the low ionic strength conditions of our optimized assay. This observation is also consistent with the previously observed requirement of a ≥12-bp region surrounding methyl-CpG sites for stable MBD binding^11^.

Next, we investigated a series of target designs with two symmetrical methylation sites: M2c-A39m-CP, M2b-A39m-CP and M2a-A39m-CP (**Fig. 4**). We hypothesized that MBD binding to M2c-A39m-CP and M2b-A39m-CP, whose methylations do not occur in a close tandem, would be less stable than to M2a-A39m-CP, which comprises two tandem internal methyl-CpG sites. As expected, M2c-A39m-CP and M2b-A39m-CP showed no and little evidence, respectively, of repeated MBD binding. In contrast, M2a-A39m-CP exhibited significant, highly repetitive binding of MBD with an *N_b+d_* of 152, a *τ_on_* of 0.7 s, and a *τ_off_* of 3.0 s (**Fig. 4**, **Supplementary Fig. 15** and **Supplementary Fig. 16**). These findings show that the presence of at least two tandem internal methylations in a dsDNA target yields significantly more stable binding than either a single internal symmetric methylation (M1a-A39m-CP) or two well-separated methylations (M2a- and M2b-A39m-CP)—consistent with the greater stabilization of MBD binding we previously observed for a stem-loop with two tandem methylated CpGs compared to a stem with only one symmetric methylation (**Fig. 3d,e**).

We further investigated two designs with three sets of symmetric methylations: M3b-A39m-CP and M3a-A39m-CP (**Fig. 4**). While both have three symmetric methyl-CpG sites, M3b-A39m-CP lacks a set of tandem methylations and includes a terminal methyl-CpG, and thus was expected to exhibit less stable MBD binding than M3a-A39m-CP, which has an internal pair of tandem methyl-CpG sites. Consistent with this expectation, and further supporting our overall model, M3b-A39m-CP exhibited evidence of MBD binding with an average *N*_b+d_ of only 152 among fewer molecules N with a lower *τ_on,median_* (though a similar *τ_off,median_*) compared to M2a-A39m-CP (**Fig. 4**). Exponential fitting yielded an estimated *τ_on_* of 0.3 s for M3b-A39m-CP, approximately half that of M2a-A39m-CP (which has one less methyl group), and a *τ_off_* of 2.5 s (**Supplementary Fig. 15** and **Supplementary Fig. 16**). An accumulation of *τ_on.median_* values close to 0.1 s was observed for M3b-A39m-CP, indicating that a significant fraction of binding events exhibits an actual *τ_on_* lower than 0.1 s. These results show that M3b-A39m-CP, despite having three methylation sites including two internal methyl-CpGs, does not exhibit as stable binding to MBD as M2a-A39m-CP, suggesting that tandem symmetrical methylations are critical to maximally stabilize MBD binding. Consistent with this interpretation, M3a-A39m-CP—which contains a set of tandem CpG methylations— yielded more detectable molecules with an average *N_b+d_* of 178, and a similar *τ_on.median_* and *τ_off.median_* to that of M2a-A39m-CP (**Fig. 4**). Exponential fitting gives a *τ_on_* of 0.5 s and a *τ_off_* 2.6 s for M3a-A39m-CP (**Supplementary Fig. 15** and **Supplementary Fig. 16**), indicating more stable MBD binding to M3a-A39m-CP than to M3b-A39m-CP.

Moreover, we examined a design with four methyl CpG sites (arranged as two sets of tandem methylations), M4a-A39m-CP (**Fig. 4**), which showed a similar number *N* of detected molecules to M2a-A39m-CP with an average *N_b+d_* of 152 of 167, a similar *τ_on.median_* of 0.7 s, and a significant decrease in *τ_off,median_*. Exponential fitting gives a *τ_on_* of 0.7 s and a *τ_off_* of 2.3 s for M4a-A39m-CP (**Supplementary Fig. 15** and **Supplementary Fig. 16**). This result further supports the notion that additional tandem symmetrical methylation sites increasingly stabilize MBD binding.

Finally, we tested a fully symmetrically methylated target strand, M7-A39m-CP (**Fig. 4**). Compared to M4a-A39m-CP, M7-A39m-CP showed a similar number *N* of detectable molecules with an average *N_b+d_* of 133, a similar *τ_on.median_* and a significant increase in *τ_off,median_*. Exponential fitting yielded a *τ_on_* of 0.7839 s and a *τ_off_* of 3.4158 s (**Supplementary Fig. 15** and **Supplementary Fig. 16**), suggesting that the additional methylation sites in M7-A39m-CP, including two internal and one terminal methyl-CpG, do not further stabilize MBD binding, but slightly decrease the binding rate. This slowing of MBD binding may reflect the shorter unmodified dsDNA regions remaining around the individual methyl-CpG sites that are needed for MBD landing^11^.

### Model of MBD binding to DNAs with different methylation sites and structural motifs

To summarize differences in binding caused by methylation states and structural motifs, we developed a generalized model for the MBD interaction with 5-methyl-CpGs (**Fig. 5**). First, a molecule of MBD binds to a DNA duplex from solution. Two main types of interaction facilitate this process: (1) non-specific electrostatic interactions between the positively charged residues of MBD and the negatively charged phosphate groups in DNA, and (2) CpG-specific interactions of *CYT* ∴ *AEG* ∨ *GUA* stair motifs (∴ denotes the cation-π interaction and ∨ denotes hydrogen bonding) regardless of methylation state^12,22^. This semi-random attachment does not guarantee MBD binding to key structural motifs or methylation sites. Therefore, subsequent to initial binding, MBD undergoes fast unbiased 1-dimensional (1D) diffusion along the DNA backbone to search for potential 5mCpG sites, which competes with dissociation from the DNA. Although not directly observed for the MBD studied here, this 1D diffusional searching was demonstrated for the MBD of chicken MBD2^23,24^ and has been suggested for that of MBD4^25^. Considering their high sequence homology, we propose that our human MBD1-derived MBD undergoes a similar 1D search before achieving stable binding. Since the diffusion coefficient of the MBD of chicken MBD2 was reported to be ∼0.09 µm^2^/s^23^, it will on average take less than 1 ms to scan by Brownian diffusion a 55-bp dsDNA like the BCAT1 target used in this study. Assuming similar rates of 1D diffusion for MBD, our 100 ms time resolution is likely too long to detect transiently bound and diffusing MBD prior to dissociation, explaining why no binding is detected for designs with few or no methylation sites.

**Fig. 5.**
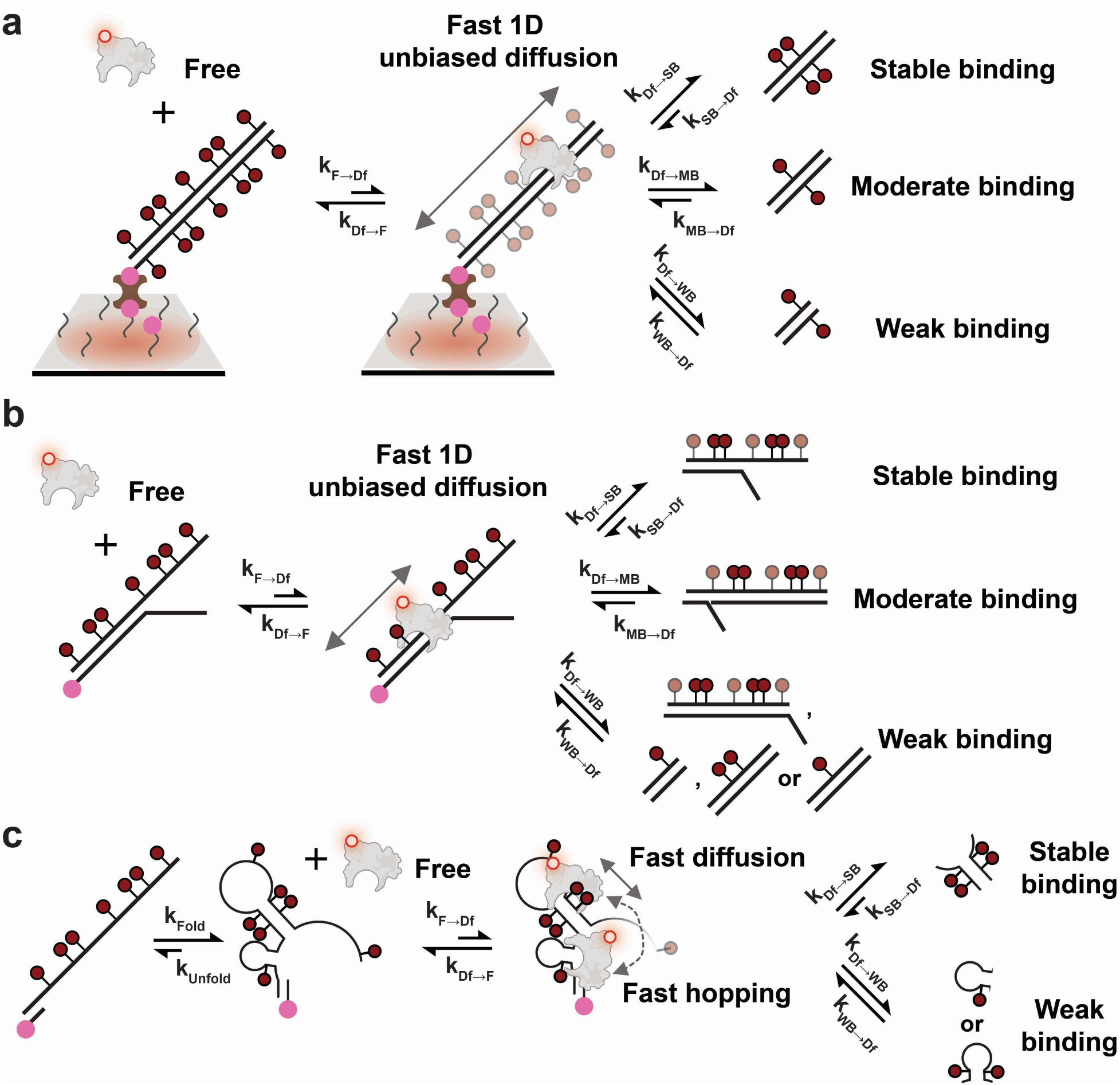
Kinetic model of human MBD1 MBD binding to DNA modulated by methylation states and structural motifs. **a** Diffusion-mediated MBD binding to symmetrically methylated dsDNA. **b** Diffusion-mediated MBD binding to bifurcating hemimethylated DNA. Semi-transparent methyl groups are not required to produce stable or moderate binding. **c** Diffusion-mediated MBD binding to methylated ssDNA.

During 1D diffusion, MBD may encounter and bind to 5mCpGs embedded within a duplex or other structural motif. Given the range of bound times observed, we classify MBD interactions into three types: stable binding, moderate binding and weak binding, based on their observed *τ_on_*. Stable binding represents any interaction with a *τ_on_* above 0.4 s, or 4-fold of our 100-ms exposure time; moderate binding is any interaction with *τ_on_* between 0.4 s and 0.1 s; and weak binding any interaction that is below our time resolution and largely or wholly undetectable in our single-molecule measurements. For simplicity in the model, we assume the association rate constants for strong, moderate, and weak binding (*T_Df→SB_*, *T_Df→WB_*, and *T_Df→MB_*) are similar, and that any differences in binding stability are primarily the result of different dissociation rate constants (*T_SB→Df_*, *T_MB→Df_*, and *T_WB→Df_*) for each type of interaction (**Fig. 5**).

As a further simplification, we posit that all binding to eligible methylation sites must be mediated by a fast 1D diffusion or hopping process, rather than binding directly from solution. We treat the direct binding or dissociation of free MBD from solution as a special case where the distance to the potential binding site is extremely short and thus a successful search becomes almost instantaneous. Similarly, direct dissociation is treated as dissociation after an extremely (infinitesimally) short diffusive search. Using this model, we can correlate our measured *τ_on_* and *τ_off_* with microscopic rate constants. Since free MBD is the fluorescence “off” state for immobilized DNA molecules, *T_F→Df_* is simply 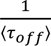; and because MBD diffuses or hops fast relative to the size of our DNAs, *T*_Df→F_, *T*_Df→SB_, *T*_Df→WB_, and *T*_Df→MB_ are assumed to be much larger than the other rate constants, causing an undetected short-lived diffusion state (**Fig. 5**). Therefore, the three types of binding classes (stable, moderate and weak) primarily contribute to the length of the fluorescence “on” state. Specifically, *τ_on_* is the weighted average of 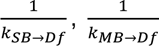, and 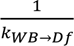 (i.e., weighted by the time-averaged probability of engaging in each class of interaction) and, in cases where stable binding exists, is dominated by 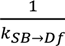 since 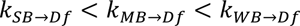.

## Discussion

In this work, we have systematically investigated the influence of the number and positioning of methylation sites and different structural motifs encountered during the cellular processes of genome transcription, replication and repair on the binding of the human MBD1 5mCpG-binding domain (MBD) using a 55-bp fragment of the BCAT1 promoter, an epigenetic biomarker for colorectal cancer, as a prototypical target. Previous structural or mechanistic studies have focused either on a single methylation site or on characterizing MBD binding to 5-methyl-CpG clusters over 1 kb in length^12,15,16,23,26–31^. In genomic studies, CpG islands generally are identified as regions over 200 bp in length that are treated as a single genetic element^32^. However, local variations of methylation patterns within 100 ∼ 200 bp have largely been ignored both in structural or mechanistic studies and in genomic studies, despite the fact that—as we have shown here—they have a strong impact on recognition by a representative methylation-sensitive protein of the MBD superfamily. We here addressed this knowledge gap by measuring the equilibrium binding kinetics of MBD to diverse methylation patterns and structural motifs. We discovered two significant properties of DNA that promote stable binding of MBD: 1) tandem symmetrically methylated CpGs in longer dsDNA and shorter stem loops formed by ssDNA; and 2) DNA fork motifs formed by single-stranded extensions in hemimethylated dsDNA. Based on our findings, we propose a mechanistic model that provides a framework for understanding the function of the MBD superfamily as epigenetic reader proteins governing the recruitment of chromatin modifying complexes, regulation of gene silencing, and prevention of epigenetic mark spread. Specifically, our results advance the mechanistic understanding of how MBDs recognize 5-methylated CpG sites in DNA and maintain the epigenetic boundaries within the human genome based on the observed limited 1D diffusion along unmethylated dsDNA.

In addition to their canonical roles in silencing transcription by recruiting histone modifiers and chromatin remodelers, MBD proteins have also been shown to mediate heterochromatin maturation at replication foci where both DNA fork and methylated ssDNA can transiently occur^33^. The affinity of MBD for hemimethylated DNA forks and methylated ssDNA might help MBD2-MBD3 or MBD1 recognize the replication fork and recruit HDACs and CAF1 to synchronize epigenetic maintenance and heterochromatin maturation^9,33^. Beyond MBD’s ability to recognize single symmetrical methyl-CpG sites, the increased binding affinity for tandem symmetrical methylation sites might be an evolutionary adaptation to increase discrimination of MBD between unmethylated and highly methylated genomic loci, which is limited in the case of isolated CpGs. For example, the K_D_ of MBD for isolated symmetrically methylated CpGs is 0.6 ∼ 0.9 µM, less than 10-fold tighter than that for a single hemimethylated CpG^16^. Although other domains of MBD proteins may provide additional affinity, considering the >5% prevalence of hemimethylated CpGs within CpG islands^34,35^ and the fact that the vast majority of all genomic CpG islands are unmethylated^36,37^, cooperative binding of tandem symmetrical methylations to MBD proteins may be essential for specifically identifying high-density methyl-CpGs while not responding to sporadic methylated CpGs in the genome.

It remains to be seen how the properties of a prototypical MBD discovered here influence the binding and other functions of full-length MBD proteins. Their other domains serve distinct functions in recruiting histone modifiers and chromatin remodelers and thus modify specifically those sites that engender tight MBD binding to affect transcriptional regulation and other biological processes. Future *in vitro* studies involving full-length MBD proteins and their co-regulators can build on the framework provided by our mechanistic model to address these questions. While our kinetic model does not detail the exact mechanism of MBD binding to DNA forks, nor fully explains all nuances in our data such as the slower binding of fully methylated symmetrical dsDNA (M7-A39m-CP in **Fig. 4**), future approaches such as MD simulations informed by our model might do so.

## Methods

### Oligonucleotides

All DNA oligonucleotides were purchased from Integrated DNA Technologies (IDT, www.idtdna.com) with standard desalting purification, unless otherwise noted. The 55 nt promoter sequence of branched-chain amino acid transaminase 1, BCAT1 was chosen as our detection target — its genomic coordinates were Chr12: 24,949,105 - 24,949,159 (genome build: UCSC Genome browser GRCh38/hg38 version)^18,19^. See **Supplementary Table 1** for descriptions of each target and their acronyms.

### Cloning, expression and purification of MBD-Halo

To generate the design for expressing MBD-Halo, a HaloTag-containing vector pBD003_mut_VCP (R155H) was gifted to us from Stephanie Moon’s lab in the Human Genetics Department at the University of Michigan. To fuse MBD1 MBD (aa 1-77) with a C-terminal HaloTag using Gibson assembly, we first PCR linearized pBD003_mut_VCP (R155H) with a pair of backbone primers: MBD-Halo-BF and MBD-Halo-BR, followed by gel purification. Insert containing MBD1 MBD (aa 1-77) was then generated by PCR amplification of Addgene plasmid # 119966 with a pair of insert primers: MBD-Halo-IF and MBD-Halo-IR, followed by gel purification. Finally, the linearized backbone and insert were mixed in a 1:2 molar ratio and ligated using Gibson assembly (NEB, Cat. # E5510S) at 50°C for 15 min. The ligation mix was then transformed into NEB 5-alpha competent cells (NEB, Cat. # C2987H) and colonies were selected using 50 µg/mL Kanamycin LB agar plate. Plasmid sequence was validated by Sanger sequencing. For overexpression and purification of MBD-Halo, its design was transformed into BL21(DE3) competent E. coli (NEB, Catalog # C2527H) and transformed cells were spread on a 50 µg/mL Kanamycin LB agar plate. Single colonies were inoculated into a 10 mL LB culture containing 50 µg/mL Kanamycin and grown overnight at 250 rpm, 37°C. The OD600 of this overnight culture was measured and a certain amount of it was further inoculated into a 100 mL TB culture with 50 µg/mL Kanamycin for large-scale expression such that the starting OD600 was exactly at 0.01. After incubating at 250 rpm, 37°C for approximately 3-4 h, its OD600 reached 0.6 and overexpression of MBD-Halo was induced by addition of 0.05 M IPTG right after cooling in ice bath. Large expression culture was further incubated at 250 rpm 20-22°C for another 16 h. After expression, E. coli culture was spun down at 5,000 g, 4°C for 20 min and cell pellets were pooled and resuspended in a lysis buffer containing 1X Base buffer (2X Base buffer contained 40 mM Tris-HCl pH 8.0, 0.2% (v/v) Tween 20, 1200 mM NaCl and 20 mM imidazole and was premixed), 1X protease inhibitor cocktail (prepared by dissolving 2 tablets of cOmplete™, EDTA-free Protease Inhibitor Cocktail, Millipore Sigma, Cat. # 11873580001, in 50 mL buffer), 1 mg/mL lysozyme (Millipore Sigma, Cat. # L6876-10G) and 5 mM freshly thawed β-mercaptoethanol. Otherwise, cell pellets would be flash-frozen and be stored at -80°C until purification. In general, 50 mL lysis buffer was used per 100 mL of TB cell culture. Cell lysis was achieved by sonication in ice water with 5 s on and 15 s off at 70% amplitude for 20 min (total time) until cell suspension became semitransparent. Subsequently, lysate was spun down at 20,000 g, 4°C for 60 min. Supernatants were further clarified through a 0.45 µm syringe filter (Millipore Sigma, Cat. # SLHV033RS) before sample application using a gravity column (Bio-Rad, Cat. # 7372512). A 2 mL Ni-NTA resin (Qiagen, Cat. # 30210) was equilibrated with 10 mL of 1X Base buffer and 5 mM β-mercaptoethanol for 10 min by constantly rotating. Following that, filtered supernatants were incubated with resin for 60 min by constantly rotating. MBD-Halo-bound resin was then slowly depositing into the gravity column and washed by 20 mL of 1X Base buffer and 5 mM β-mercaptoethanol. A series of step gradient of elution buffers were used containing: 20 mM imidazole, 50 mM imidazole, 80 mM imidazole, 100 mM imidazole, 150 mM imidazole and 200 mM imidazole in 1X Base buffer with 5 mM β-mercaptoethanol. Each gradient step was 10 mL and each fraction of eluate was around 5 mL. Fractions were loaded on denaturing PAGE (NuPAGE 4-12% Bis-Tris protein gels, 1.0 mm, 17-well, Fisher Scientific, Cat. # NP0329BOX) to examine protein purity. Combined pure MBD-Halo were concentrated and buffer-exchanged in a storage buffer containing 20 mM Tris-HCl pH 8.0, 0.1% (v/v) Tween 20, 10% (v/v) glycerol, 300 mM NaCl and 5 mM β-mercaptoethanol. Finally, MBD-Halo was aliquoted, flash-frozen in liquid nitrogen and stored at -80°C. All steps following cell harvest were at 4°C.

### Labeling of MBD-Halo

To label MBD-Halo with HaloTag ligand Alexa Fluor 660 (AF660, Promega, Cat. # G8471), 1 µM of MBD-Halo was mixed with 5 µM AF660 in a reaction buffer of 300 mM Sodium Phosphate pH 7.4, 0.1% (v/v) Tween 20, 10% (v/v) glycerol and 5 mM β-mercaptoethanol for 1 h in dark at 4°C. Free AF660 was removed using 10K MWCO centrifugal filter (Millipore Sigma, Cat. # UFC501024) at 12,500 g, 4°C until all free dyes were removed (examined by denaturing PAGE). Labeling efficiency was estimated to be close to 100% by gel imaging (data not shown here). Finally, 50% (v/v) glycerol stock was prepared, aliquoted and stored at -20°C.

### Electrophoresis mobility shift assay (EMSA)

Polyacrylamide was used for EMSA to examine methyl-CpG binding activity of MBD-Halo and MBD-Halo-AF660. 5% native PAGE were prepared in 50 mM Tris Acetate pH 7.5. 100 nM dsDNA substrates and proteins at different molar ratios were incubated in the binding buffer of 10% glycerol, 50 mM Tris-HCl pH 8.0 at room temperature for 2 h. Electrophoresis was running in 50 mM Tris Acetate pH 7.5 at 4°C with approximately 15 V/cm for 3 h. Gel was stained by SYBR Gold and visualized using Cy2 fluorescence on Typhoon (Cytiva, Amersham™ Typhoon™ Biomolecular Imager).

### Single-molecule fluorescence microscopy

Sample cells made of cut P20 pipette tips were attached to glass coverslips passivated with a 1:100 mixture of biotin-PEG and mPEG. A detailed protocol of slide preparations is discussed elsewhere^38,38–40^. Sample cells were first washed with T50 buffer (10 mM Tris-HCl pH 8.0 at 25°C, 50 mM NaCl) and then incubated with 40 µl 0.25 mg/mL streptavidin in T50 buffer for 10 min. Following wash with 1X PBS for 3 times, 100 nM capture probe in 1X PBS was prepared by heating at 85°C for 5 min in a metal bath, annealed at 37°C for 5 min in a water bath, cooled down to room temperature, and then was added to the sample well for 10-min incubation. Following wash with 4X PBS for 3 times, a mixture of target components was prepared in a PCR tube that contained 10 nM antisense strand and 10 pM targets in 4X PBS / 2 µM poly-T oligodeoxyribonucleotide (dT10) carriers. PCR tubes that contained target components including targets were then heated at 80°C for 3 min, annealed at 64°C for 5 min, subsequently 57°C for 5 min and cooled down at 38°C for another 5 min and finally held at 22°C. This target assembly process was performed in a thermocycler. The target design that was properly assembled was added to the sample cell and then incubated for 1 h at room temperature. After target capture, sample cells were washed 3 times with 4X PBS followed by one-time wash of 50 mM Tris-HCl pH 8.0. 50 ∼ 100 µl imaging buffer containing the desired concentration of MBD-Halo-AF660 in the presence of an oxygen scavenger system (OSS) — 1 mM Trolox, 5 mM 3,4-dihydroxybenzoate (PCA), 50 nM protocatechuate dioxygenase (PCD) — was added and then imaged by objective-TIRF microscopy. 1 µM PCD stock was prepared in 100 mM Tris-HCl pH 8.0, 50 mM KCl, 1 mM EDTA, 50% glycerol; 100 mM PCA was dissolved in water and titrated with 5 M KOH to a pH of 8.3; Trolox was dissolved in water and titrated with 5 M KOH to a pH around 10-11. All three components were stored at -20°C prior to use.

### Image acquisition

All single-molecule experiments were performed on the Oxford Nanoimager (ONI), a compact benchtop microscope capable of objective-type TIRF (See https://oni.bio/nanoimager/ for spec sheet regarding camera, illumination and objective). A 100X 1.4NA oil-immersion objective was installed on ONI together with a built-in Z-lock control module for autofocus. Since the built-in temperature control system on ONI could not keep imaging temperature below 25°C, to avoid overheating due to turning laser on for too long, we attached the outer box of ONI to a metal clamp where circulating cold water from water bath can come through. To maintain an imaging temperature of 22°C, water bath was kept at 16°C. For recording AF660 fluorescence emission with optimal signal-to-noise ratio (S/N), samples were excited at 640 nm with 20% laser power (approximately 30 mW) at an illumination angle of 54.0° ∼ 54.3° (note that this “illumination angle” shown on ONI was not actually the incident angle. The relationship between illumination angle and incident angle was not clear to us.). The signal integration time (exposure time) per frame was 100 ms unless otherwise noted, movies of 5 min were collected per field of view (FOV).

### Processing and analysis of objective-TIRF data

A set of custom MATLAB codes was used to identify spots with significant intensity fluctuations within each FOV, generate intensity-versus-time traces at each spot, fit these traces with a two-state hidden Markov model (HMM) to generate idealized traces, and eventually identify and characterize transitions with idealized traces. (**Supplementary Table 2**). Detailed discussions of the data analysis pipeline are published elsewhere^38–42^.

## Acknowledgments

We thank Andreas Schmidt, now at Illumina, for feedback and Stephanie Moon from the Department of Human Genetics at the University of Michigan for providing pBD003_mut_VCP (R155H). This work was supported by NIH grants R21 CA225493 (N.G.W. and M.T.) and R35 GM131922 (N.G.W.).

## Author contributions

L.D. and N.G.W. conceived and designed the study. L.D. performed experiments and analyzed the data. L.D., A.J.-B. and N.G.W. wrote the manuscript. N.G.W. and A.J.-B. supervised the study.

## Inclusion and diversity

We support inclusive, diverse, and equitable conduct of research.

