## Supplementary Information for "Mechanistic model for epigenetic maintenance by methyl-CpG-binding domain proteins"

The manuscript was written based on contributions from all authors.

All authors have given approval to the final version of the manuscript.

**Supplementary Table 1.** DNA oligonucleotide names, sequences, and descriptions. All sequences are listed 5′-to-3′.

| Name | Sequences | Description |
|---|---|---|
| | **Cloning** | |
| Halo-hMBD-BF | GGAGGTGGAAGCGGTGAA | Forward primer for linearization of backbone plasmid pBD003_mut_VCP (R155H) for constructing Halo-hMBD, directly purchased from IDT |
| Halo-hMBD-BR | ATGTATATCTCCTTCTTAAAGTTAA | Reverse primer for linearization of backbone plasmid pBD003_mut_VCP (R155H) for constructing Halo-hMBD, directly purchased from IDT |
| Halo-hMBD-IF | GAAGGAGATATACATATGGCTGAGGACTGGCTGG | Forward primer for PCR amplification of insert hMBD1 MBD from Addgene plasmid #119966 for constructing 81 Halo-hMBD, directly purchased from IDT |
| Halo-hMBD-IR | ACCGCTTCCACCTCCATGGGCCTTGGGGGCTGG | Reverse primer for PCR amplification of insert hMBD1 MBD from Addgene plasmid #119966 for constructing Halo-hMBD, directly purchased from IDT |
| | **Target** | |
| M7 | GTCTTCCTGCTGATGCAATC/iMe-dC/GCTAGGT/iMe-dC/G/iMe-dC/GAGTCTC/iMe-dC/GC/iMe-dC/G/iMe-dC/GAGAGGGC/iMe-dC/GG | Fully Methylated 55 nt BCAT1 promoter forward strand, with 7 methyl-CpGs, directly purchased from IDT |
| M4b | GTCTTCCTGCTGATGCAATCCGCTAGGT /iMe-dC/ G /iMe-dC/GAGTCTC/iMe-dC/GC/iMe-dC/GCGAGAGGGCCGG | Methylated 55 nt BCAT1 promoter forward strand, with 4 methyl-CpGs, directly purchased from IDT |
| M4a | GTCTTCCTGCTGATGCAATCCGCTAGGT /iMe-dC/ G /iMe-dC/ GAGTCTCCGC /iMe-dC/ G /iMe-dC/ GAGAGGGCCGG | Methylated 55 nt BCAT1 promoter forward strand, with 4 methyl-CpGs, directly purchased from IDT |
| M3b | GTCTTCCTGCTGATGCAATCCGCTAGGTCGCGAGTCTC/iMe-dC/GC/iMe-dC/GCGAGAGGGC/iMe-dC/GG | Methylated 55 nt BCAT1 promoter forward strand, with 3 methyl-CpGs, directly purchased from IDT |
| M3a | GTCTTCCTGCTGATGCAATCCGCTAGGTCGCGAGTCTC/iMe-dC/GC/iMe-dC/G/iMe-dC/GAGAGGGCCGG | Methylated 55 nt BCAT1 promoter forward strand, with 3 methyl-CpGs, directly purchased from IDT |

| | | |
|---|---|---|
| M2c | GTCTTCCTGCTGATGCAATCCGCTAGGTCGCGAGTCTCCGC/iMe-dC/GCGAGAGGGC/iMe-dC/GG | Methylated 55 nt BCAT1 promoter forward strand, with 2 methyl-CpGs, directly purchased from IDT |
| M2b | GTCTTCCTGCTGATGCAATCCGCTAGGTCGCGAGTCTC/iMe-dC/GCCGCGAGAGGGC/iMe-dC/GG | Methylated 55 nt BCAT1 promoter forward strand, with 2 methyl-CpGs, directly purchased from IDT |
| M2a | GTCTTCCTGCTGATGCAATCCGCTAGGT/iMe-dC/G/iMe-dC/GAGTCTCCGCCGCGAGAGGGCCGG | Methylated 55 nt BCAT1 promoter forward strand, with 2 methyl-CpGs, directly purchased from IDT |
| M1b | GTCTTCCTGCTGATGCAATCCGCTAGGTCGCGAGTCTCCGCCGCGAGAGGGC/iMe-dC/GG | Methylated 55 nt BCAT1 promoter forward strand, with single methyl-CpGs, directly purchased from IDT |
| M1a | GTCTTCCTGCTGATGCAATC/iMe-dC/GCTAGGTCGCGAGTCTCCGCCGCGAGAGGGCCGG | Methylated 55 nt BCAT1 promoter forward strand, with single methyl-CpGs, directly purchased from IDT |
| M0 | GTCTTCCTGCTGATGCAATCCGCTAGGTCGCGAGTCTCCGCCGCGAGAGGGCCGG | Unmethylated 55 nt BCAT1 promoter forward strand, directly purchased from IDT |
| | **Auxiliary probes** | |
| A39m | C/iMe-dC/GGCCCTCT/iMe-dC/G/iMe-dC/GG/iMe-dC/GGAGACT/iMe-dC/G/iMe-dC/GACCTAG/iMe-dC/GGATT | Fully methylated auxiliary probe, 39 nt in total, fully complementary to target sequence, with no "branch" motif, with 7 methyl-CpGs, directly purchased from IDT |
| A39O | CTTATCTGTTCCGGCCCTCTCGCGGCGGAGACTCGCGACCTAGCGGATT | 49 nt auxiliary probe, with 39 nt complementary to target sequence, with a 10 nt overhang sequence "CTTATCTGTT", directly purchased from IDT |
| A39 | CCGGCCCTCTCGCGGCGGAGACTCGCGACCTAGCGGATT | 39 nt auxiliary probe, fully complementary to target sequence, directly purchased from IDT |
| A30 | TCGCGGCGGAGACTCGCGACCTAGCGGAT | 30 nt auxiliary probe, fully complementary to target sequence, directly purchased from IDT |
| A25 | GCGGAGACTCGCGACCTAGCGGATT | 25 nt auxiliary probe, fully complementary to target sequence, directly purchased from IDT |
| A17 | TCGCGACCTAGCGGATT | 17 nt auxiliary probe, fully complementary to target |

| | | |
|---|---|---|
| | | sequence, directly purchased from IDT |
| A17O | CTTATCTGTTTCGCGACCTAGCGGATT | 27 nt auxiliary probe, with 17 nt complementary to target sequence, with a 10 nt overhang sequence "CTTATCTGTT", directly purchased from IDT |
| | **Capture probes** | |
| CPO | /5Biosg/ATAATTAATAGCATCAGCAGGAAGAC | 26 nt 5' biotinylated capture probe, with 16 nt complementary to target sequence, with a 10 nt overhang sequence "ATAATTAATA", directly purchased from IDT |
| CP | GCATCAGCAGGAAGAC/3BioTEG/ | 16 nt 3' biotinylated capture probe, fully complementary to target sequence, directly purchased from IDT |
| | **Others** | |
| M7_rev | C/iMe-dC/GGCCCTCT/iMe-dC/G/iMe-dC/GG/iMe-dC/GGAGACT/iMe-dC/G/iMe-dC/GACCTAG/iMe-dC/GGATTGCATCAGCAGGAAGAC | Fully Methylated 55 nt BCAT1 promoter reverse strand, with 7 methyl-CpGs, fully complementary to M7, directly purchased from IDT |
| M0_rev | CCGGCCCTCTCGCGGCGGAGACTCGCGACCTAGCGGATTGCATCAGCAGGAAGAC | Unmethylated 55 nt BCAT1 promoter reverse strand, fully complementary to M7, directly purchased from IDT |

**Supplementary Table 2.** Parameter sets for trace generation and analysis.

| Trace Generation Parameters | |
|---|---|
| use fluctuation map? | 2 ('$N_{b+d}$ map') |
| Stdfactor | 5 |
| start frame | 1 |
| end frame | 3000 |
| edgePx | 20 |
| Percentilecut | 0.95 |
| ROI size (pixels) | 5 |

| Trace Fitting Parameters | |
|---|---|
| start frame | 1 |
| end frame | 3000 |
| exposure time (s) | 0.1 |
| Smoothframes | 1 |
| remove_single_frame_events | FALSE |
| Ithresh | 1000 |
| SNthresh | 2 |

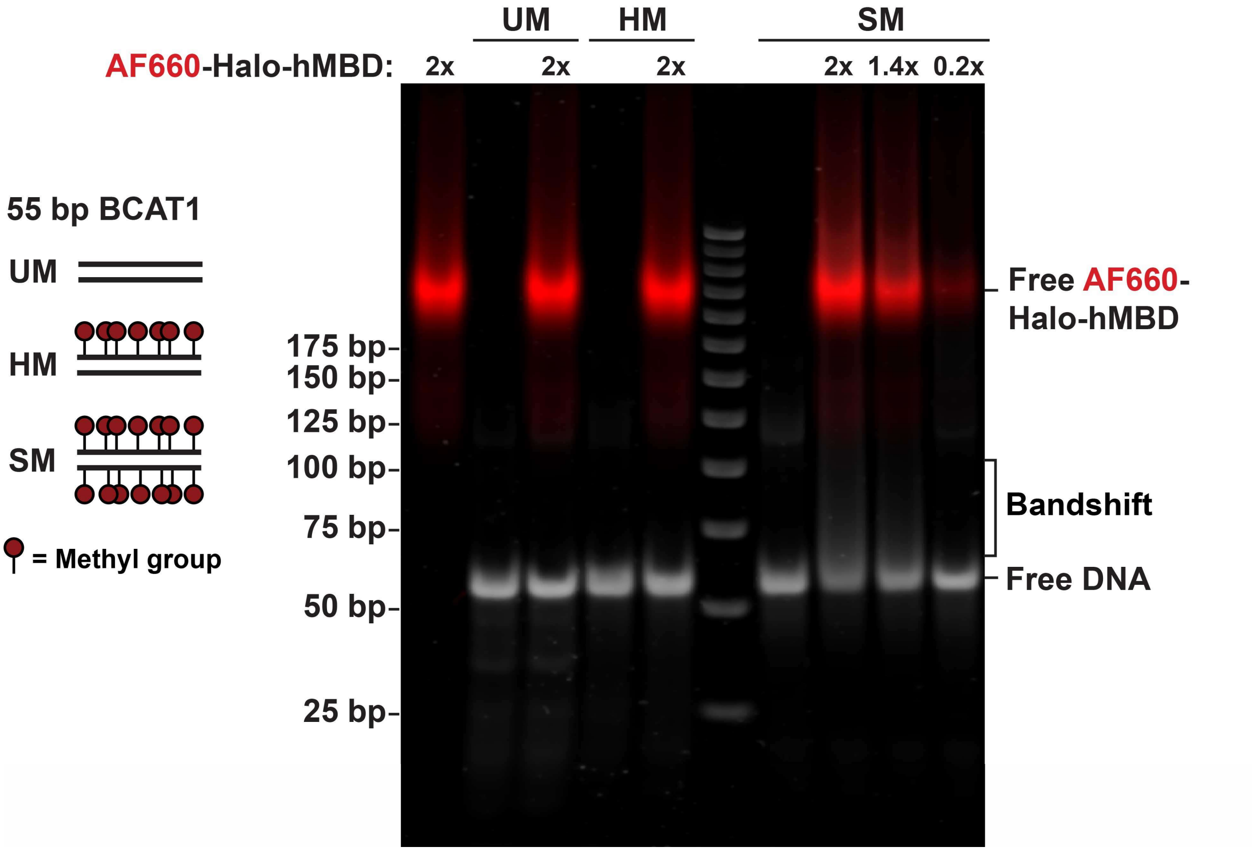


**Supplementary Fig. 1. | EMSA of AF660-Halo-hMBD binding to three types of 55 bp BCAT1 promoter substrates.** SM (symmetrically methylated DNA), HM (hemimethylated DNA), and UM (unmethylated DNA). AF660-Halo-hMBD was mixed with 100 nM DNAs at different molar ratios in 10% glycerol, 50 mM Tris-HCl pH 8.0 at room temperature in dark for 2 h. 5% PAGE was prepared in 50 mM Tris Acetate pH 7.5. Electrophoresis was running in 50 mM Tris Acetate pH 7.5 at 4°C with approximately 15 V/cm for 3 h. Gel was stained with SYBR Gold and visualized using both Cy5 and Cy2 fluorescence on Typhoon Biomolecular Imager.

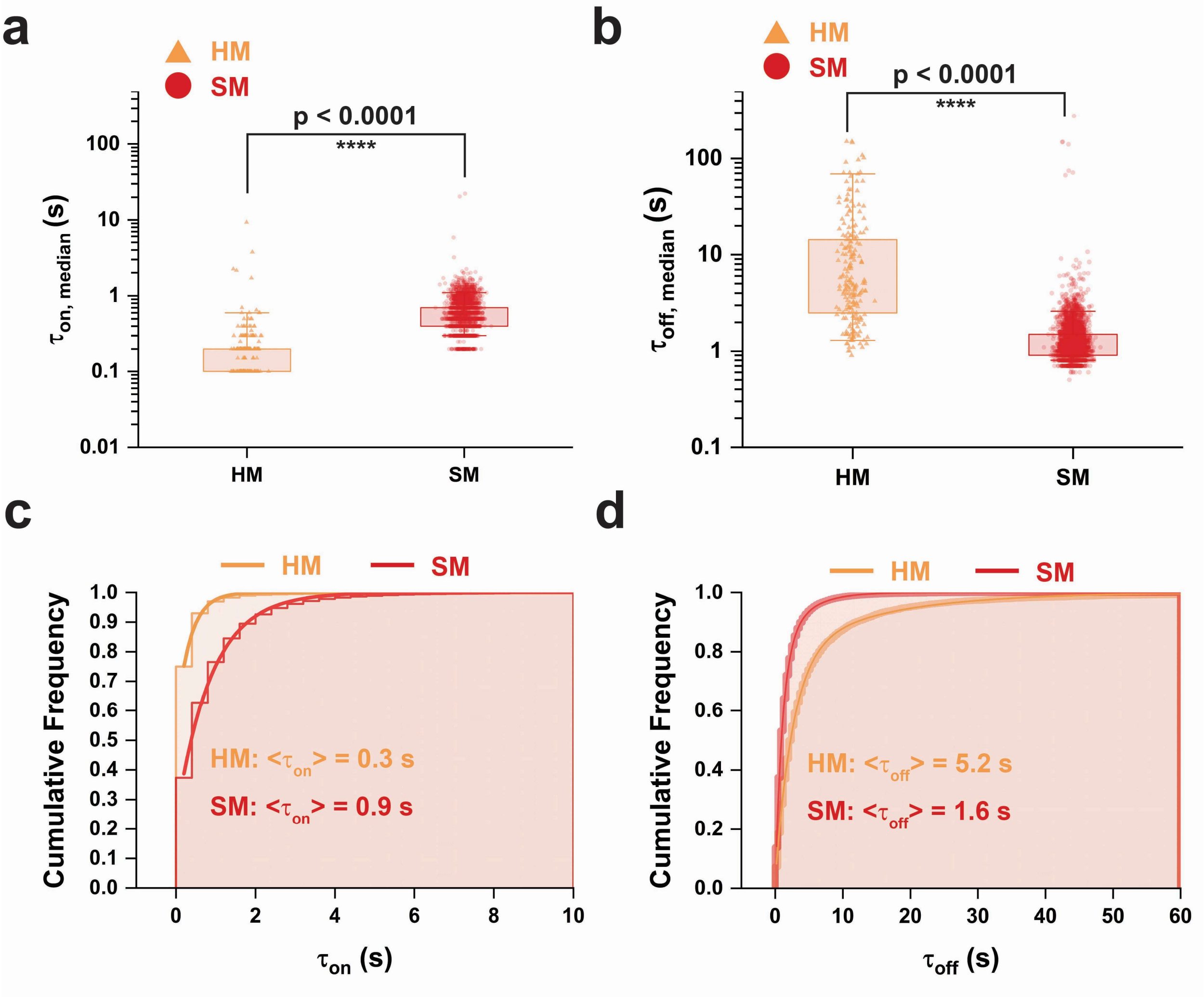


**Supplementary Fig. 2 | Dwell time comparison between SM and HM in Fig. 1. a,b** Comparison of $\tau_{on,median}$ and $\tau_{off,median}$ distributions respectively between SM and HM. Boxes are drawn from Q1 to Q3 with whiskers from 5% percentile to 95% percentile. P-values are assessed using single-tailed unpaired t-test. **c,d** Cumulative dwell time distributions and their exponential fittings of $\tau_{on}$ and $\tau_{off}$ of individual events of detected molecules. Step horizontal lines are cumulative frequency counts and solid curves are fitting curves. In panel c, mean bound time, $\langle\tau_{on}\rangle$, is calculated by fitting a single-exponential decay function. In panel d, mean unbound time, $\langle\tau_{off}\rangle$, is calculated by fitting a double-exponential decay function and taking the weighted average of individual components.

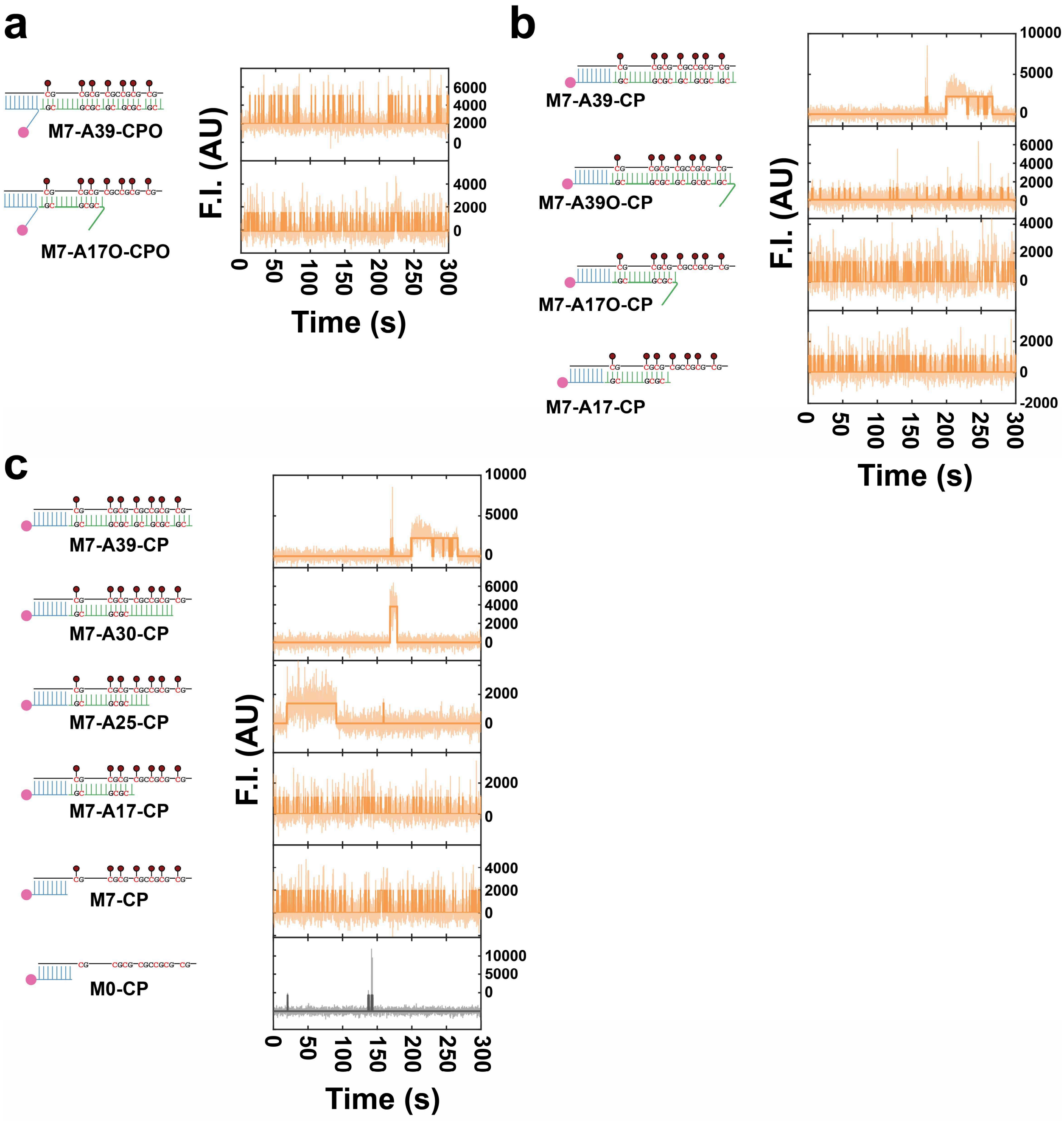


**Supplementary Fig. 3 | Representative intensity-time traces of constructs in Fig. 2.** Semi-transparent lines in the background are raw traces and solid lines are idealized traces by hidden Markov modeling. F.I., fluorescence intensity; AU, arbitrary unit.

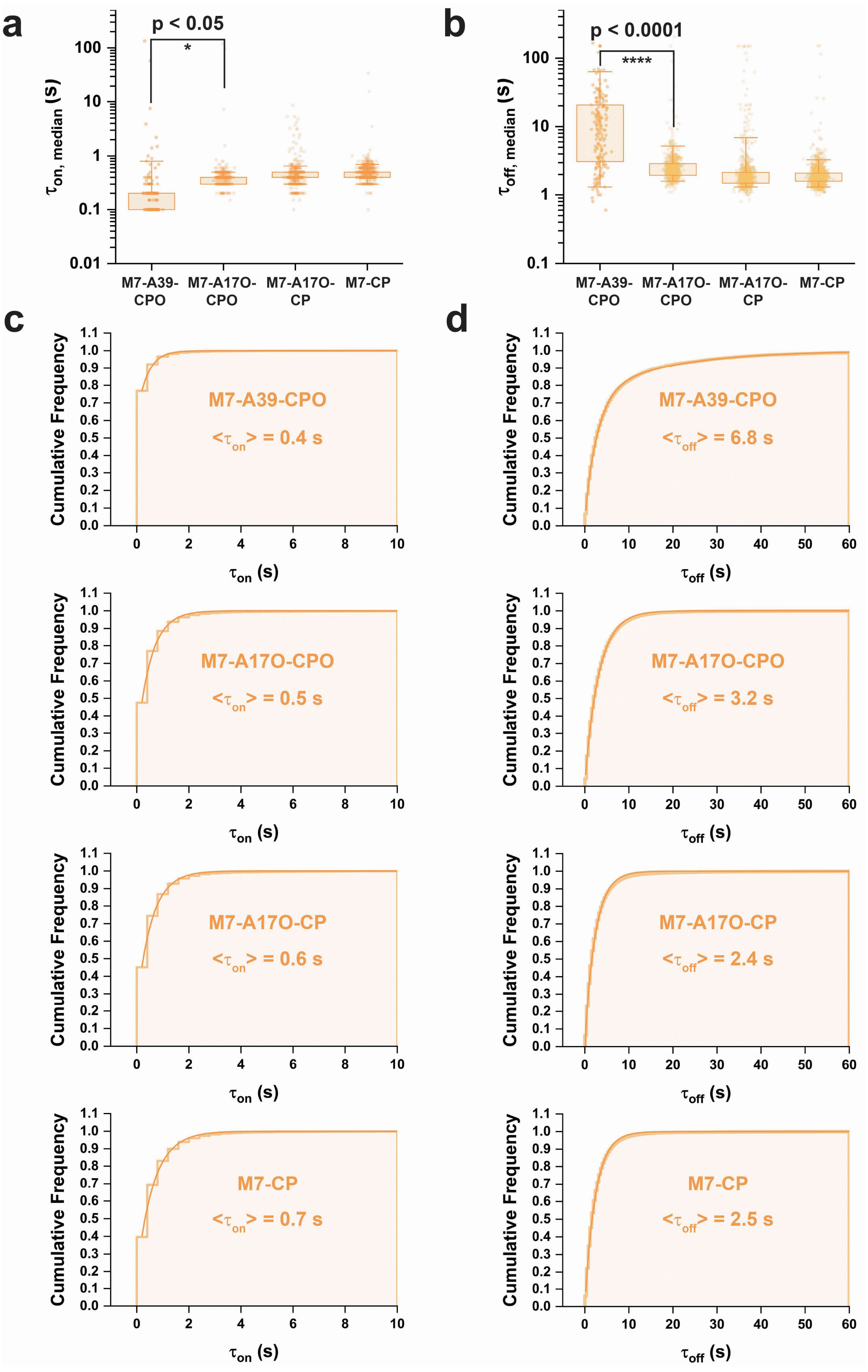


**Supplementary Fig. 4 | Dwell time comparison and mean dwell time calculations of M7-A39-CPO, M7-A17O-CPO, M7-A17O-CP and M7-CP. a,b** Boxplots of $\tau_{on,median}$ and $\tau_{off,median}$

distributions respectively. Boxes are drawn from Q1 to Q3 with whiskers from 5% percentile to 95% percentile. P-values are assessed using single-tailed unpaired t-test. **c,d** Cumulative dwell time distributions and their exponential fittings of $\tau_{on}$ and $\tau_{off}$ of individual events of detected molecules. Step horizontal lines are cumulative frequency counts and solid curves are fitting curves. Mean dwell times, $\langle\tau_{on}\rangle$ and $\langle\tau_{off}\rangle$, are calculated by fitting a single-exponential decay function except that $\langle\tau_{off}\rangle$ of M7-CP is calculated by fitting a double-exponential decay function and taking the weighted average of individual components.

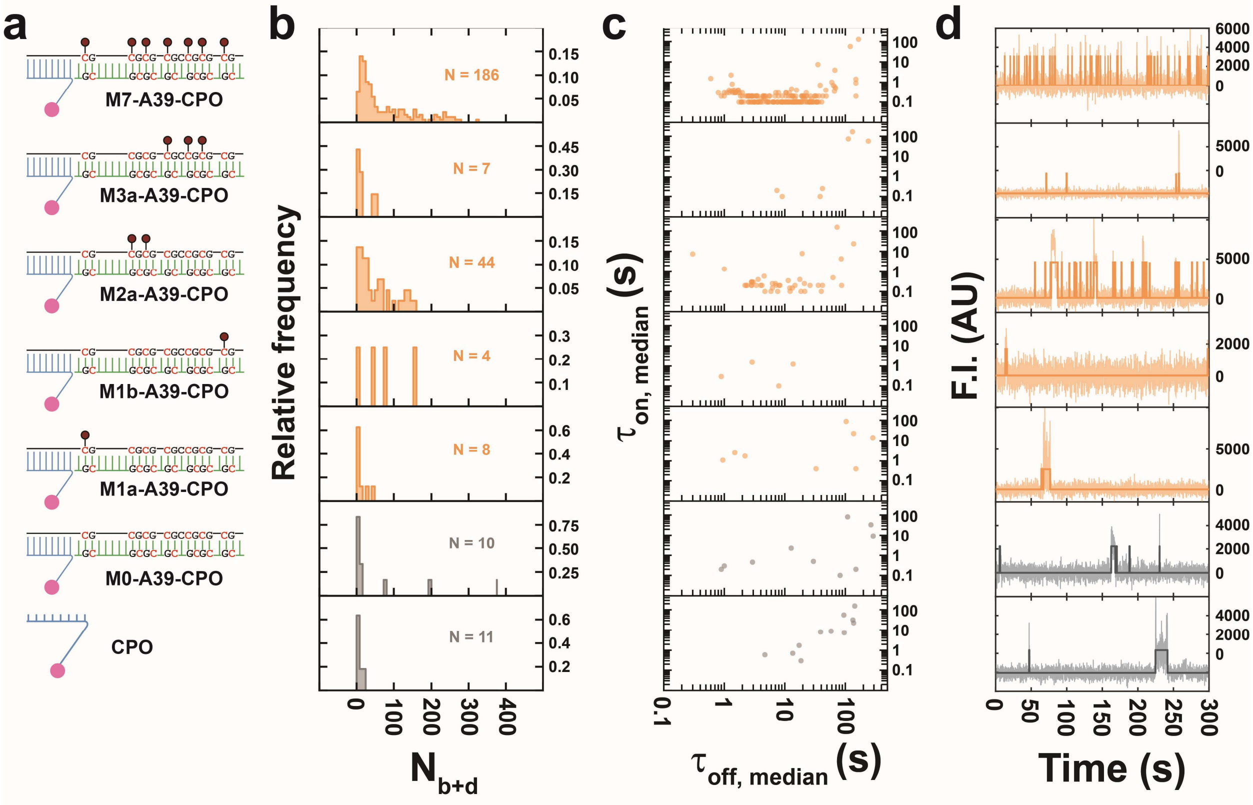


**Supplementary Fig. 5 | Binding kinetics of single internal overhang-containing constructs modulated by clustering of methyl-CpG sites. a** Constructs of M7-A39-CPO, M3a-A39-CPO, M2a-A39-CPO, M1b-A39-CPO, M1a-A39-CPO, M0-A39-CPO and CPO. **b** $N_{b+d}$ distributions. N, the number of detected molecules of one field of view (FOV). **c** Median dwell time distributions. **d** Representative intensity-time traces. Semi-transparent lines in the background are raw traces and solid lines are idealized traces by hidden Markov modeling. F.I., fluorescence intensity; AU, arbitrary unit.

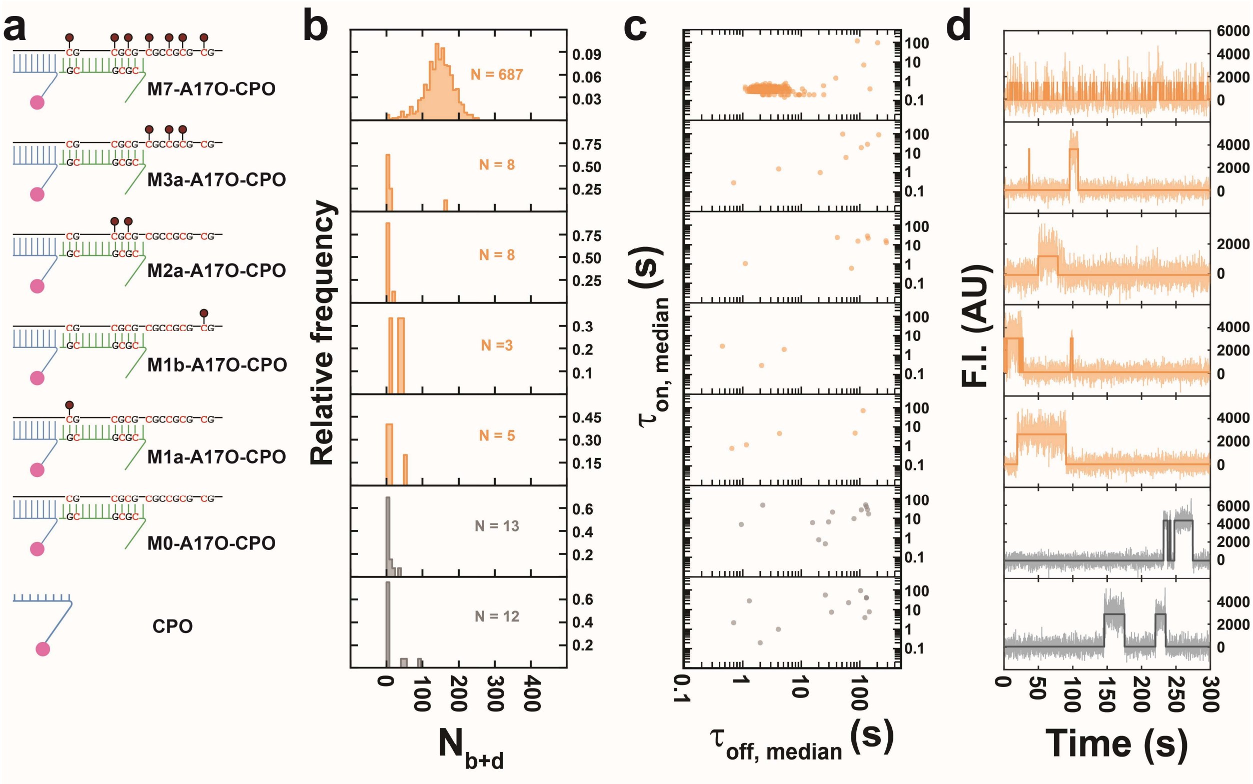


**Supplementary Fig. 6 | Binding kinetics of double overhang-containing constructs modulated by clustering of methyl-CpG sites.** **a** Constructs of M7-A17O-CPO, M3a-A17O-CPO, M2a- A17O-CPO, M1b- A17O-CPO, M1a-A17O-CPO, M0-A17O-CPO and CPO. **b** $N_{b+d}$ distributions. N, the number of detected molecules of one field of view (FOV). **c** Median dwell time distributions. **d** Representative intensity-time traces. Semi-transparent lines in the background are raw traces and solid lines are idealized traces by hidden Markov modeling. F.I., fluorescence intensity; AU, arbitrary unit.

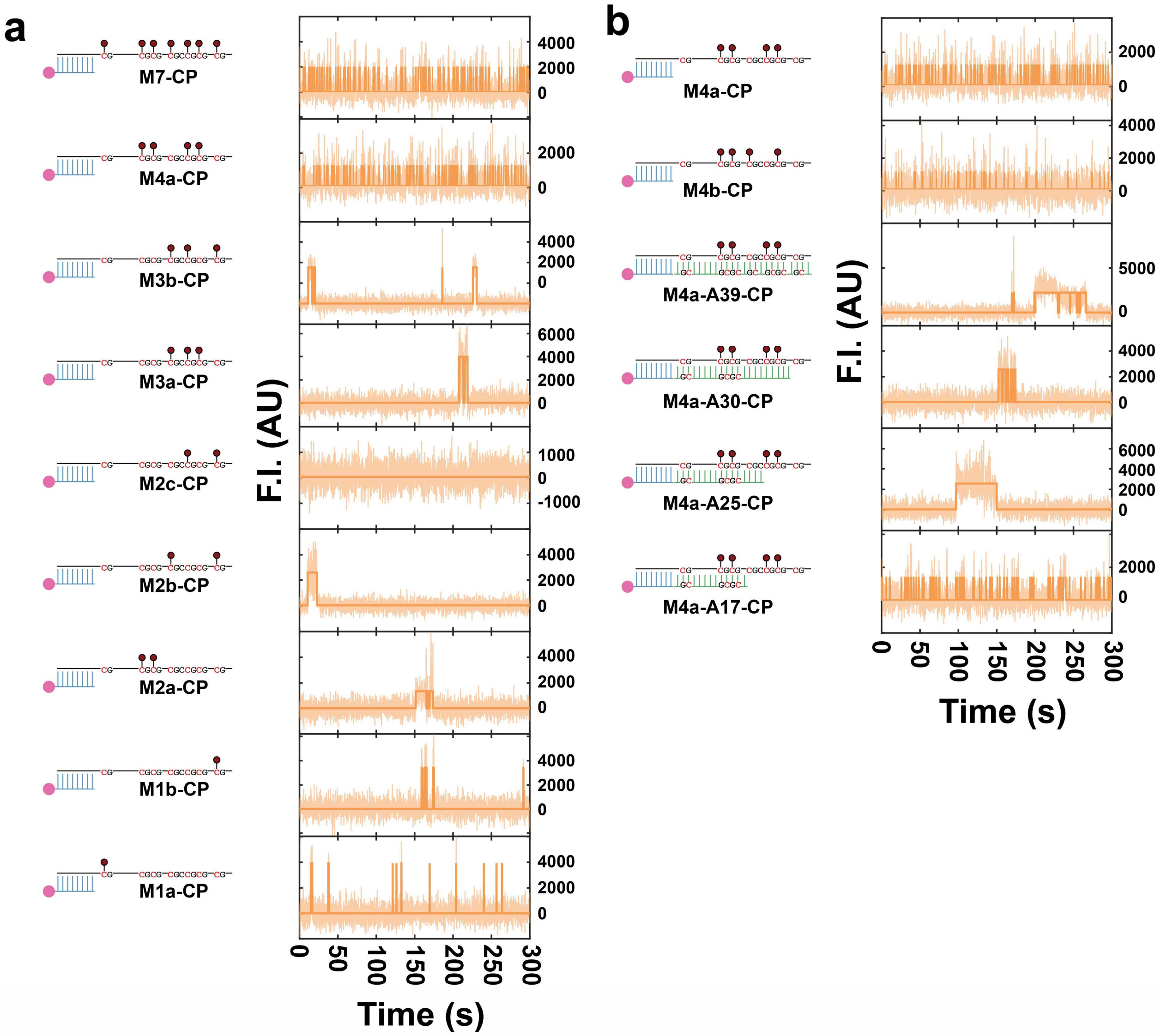


**Supplementary Fig. 7 | Representative intensity-time traces of constructs in Fig. 3.** Semi-transparent lines in the background are raw traces and solid lines are idealized traces by hidden Markov modeling. F.I., fluorescence intensity; AU, arbitrary unit.

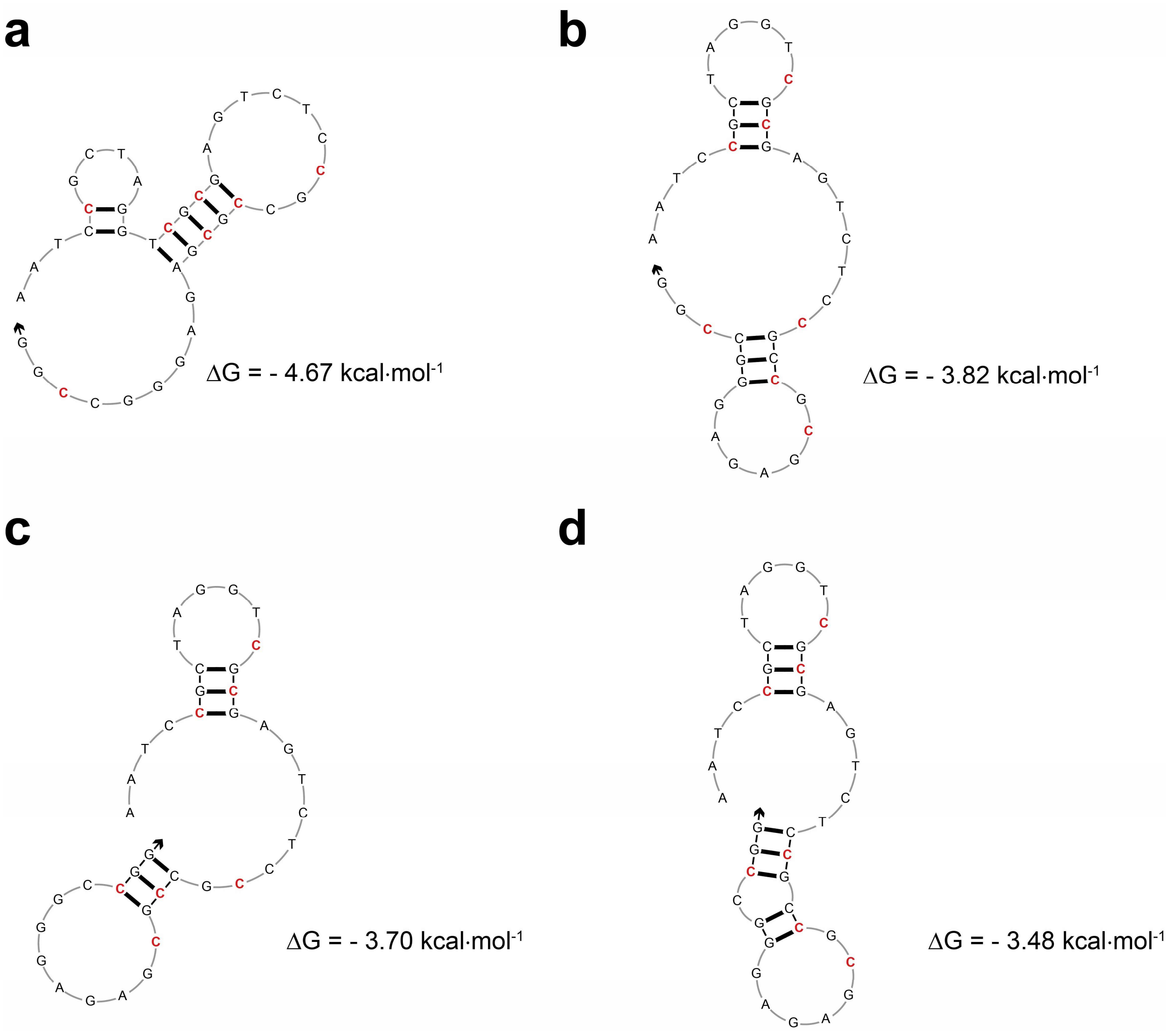


**Supplementary Fig. 8 | Predicted secondary structures and calculated free energy by IDT.** The top 4 structures are shown here. Input parameters for "HAIRPIN" prediction: Sequence, AATC**CG**CTAGGT**CGCG**AGTCTC**CG**C**CGCG**AGAGGGC**CG**G; Target type, DNA; Oligo Conc, 0.01 μM; $Na^+$ Conc, 20 mM; $Mg^{2+}$ Conc, 0 mM; dNTPs Conc, 0 mM; Suboptimality, 50%; Sequence type, Linear; Temperature, 22 °C. Max Foldings, 20; Start position, 0; Stop position, 0.

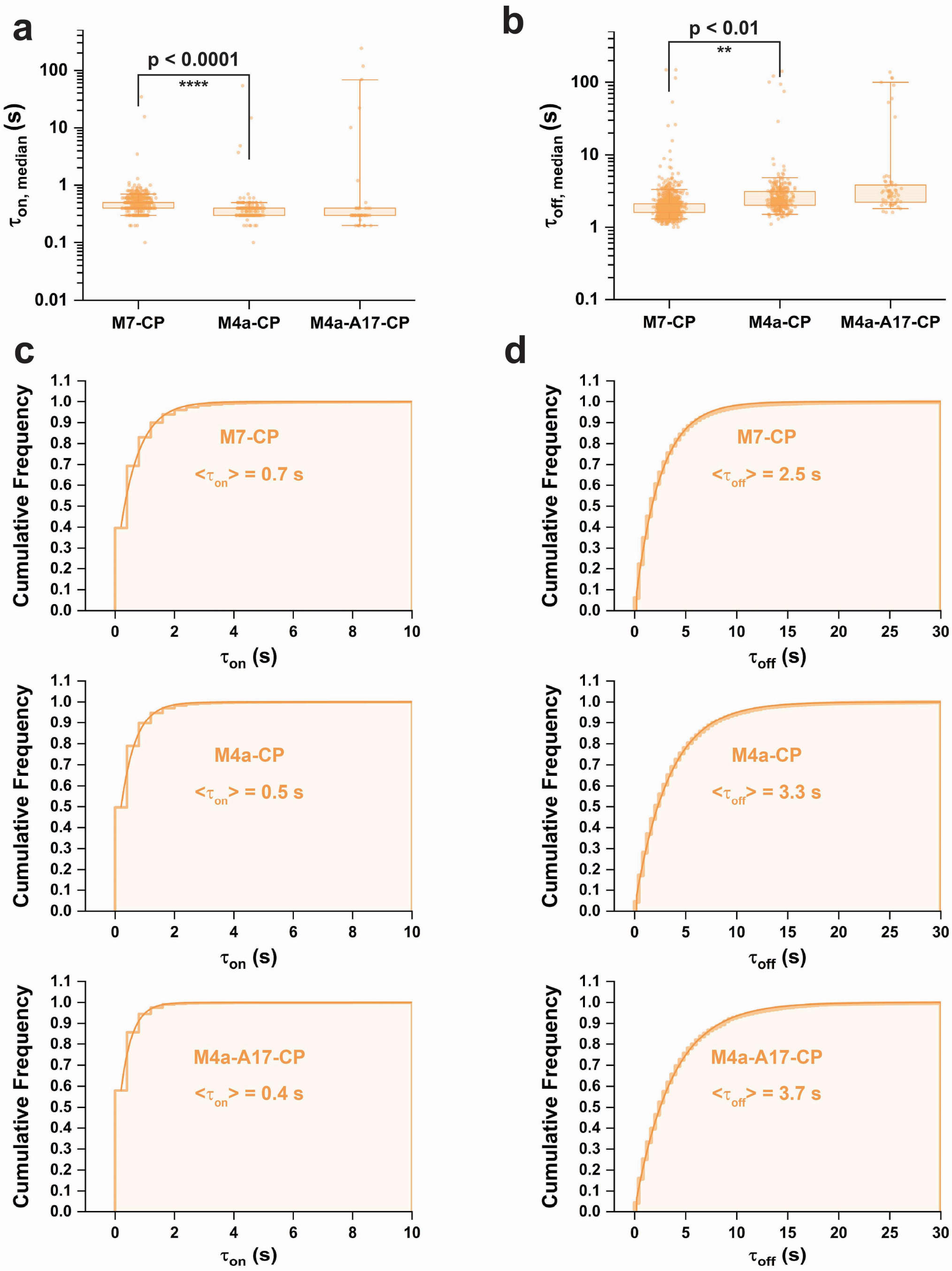


**Supplementary Fig. 9 | Dwell time comparison and mean dwell time calculations of M7-CP, M4a-CP and M4a-A17-CP. a,b** Boxplots of $\tau_{on,median}$ and $\tau_{off,median}$ distributions respectively. Boxes are drawn from Q1 to Q3 with whiskers from 5% percentile to 95% percentile. For hypothesis testing, the datasets of M4a-CP are 96% winsorized to tolerate interference of outliers. P-values are assessed using single-tailed unpaired t-test. **c,d** Cumulative dwell time distributions and their exponential fittings of $\tau_{on}$ and $\tau_{off}$ of individual events of detected molecules. Step horizontal lines are cumulative frequency counts and solid curves are fitting curves. Mean dwell times, $\langle\tau_{on}\rangle$ and $\langle\tau_{off}\rangle$, are calculated by fitting a single-exponential decay function.

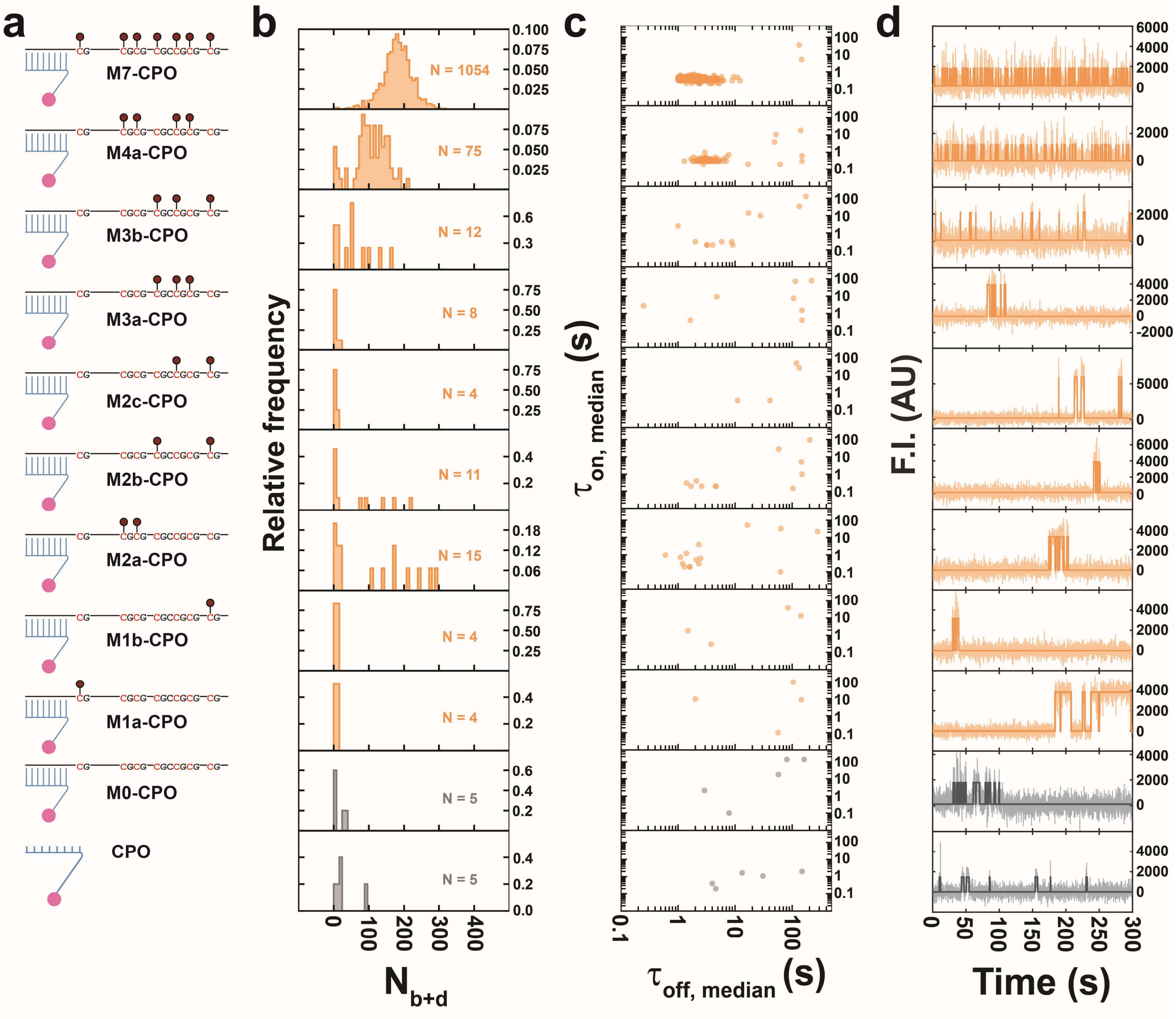


**Supplementary Fig. 10 | Interplay between overhangs and secondary structure. a** Constructs of M7-CPO, M4a-CPO, M3b-CPO, M3a-CPO, M2c-CPO, M2b-CPO, M2a-CPO, M1b-CPO and M1a-CPO, M0-CPO and CPO. **b** $N_{b+d}$ distributions. N, the number of detected molecules of one field of view (FOV). **c** Median dwell time distributions. **d** Representative intensity-time traces. Semi-transparent lines in the background are raw traces and solid lines are idealized traces by hidden Markov modeling. F.I., fluorescence intensity; AU, arbitrary unit.

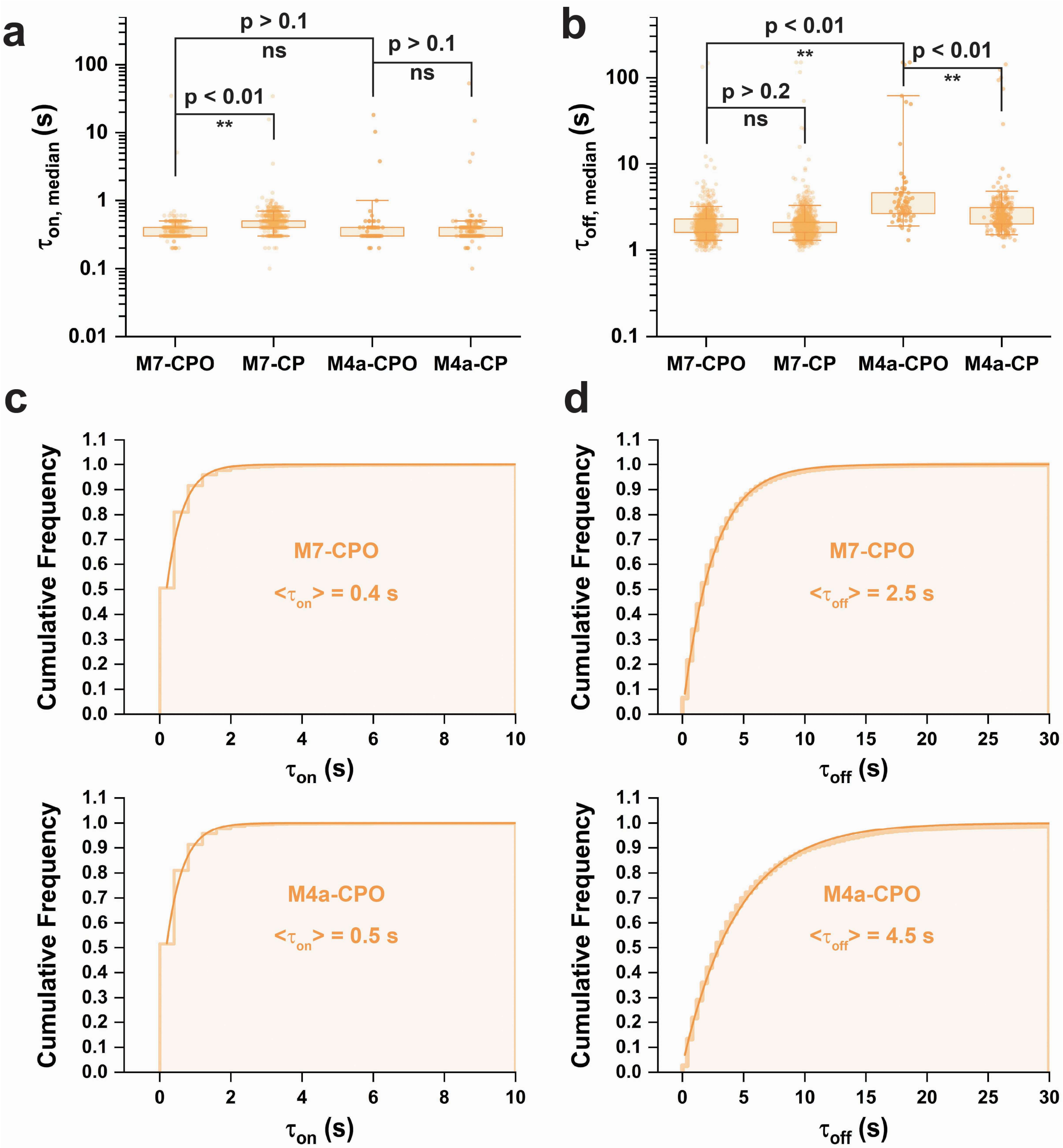


**Supplementary Fig. 11 | Dwell time comparison and mean dwell time calculations of M7-CPO, M7-CP, M4a-CPO and M4a-CP. a,b** Boxplots of $\tau_{on,median}$ and $\tau_{off,median}$ distributions respectively. Boxes are drawn from Q1 to Q3 with whiskers from 5% percentile to 95% percentile. For hypothesis testing, the datasets of M4a-CPO and M4a-CP are 96% winsorized to tolerate interference of outliers. P-values smaller than 0.05 are assessed using single-tailed unpaired t-test and P-values higher than 0.05 are assessed using two-tailed unpaired t-test. **c,d** Cumulative dwell time distributions and their exponential fittings of $\tau_{on}$ and $\tau_{off}$ of individual events of detected molecules. Step horizontal lines are cumulative frequency counts and solid curves are fitting curves. Mean dwell times, $\langle\tau_{on}\rangle$ and $\langle\tau_{off}\rangle$, are calculated by fitting a single-exponential decay function.

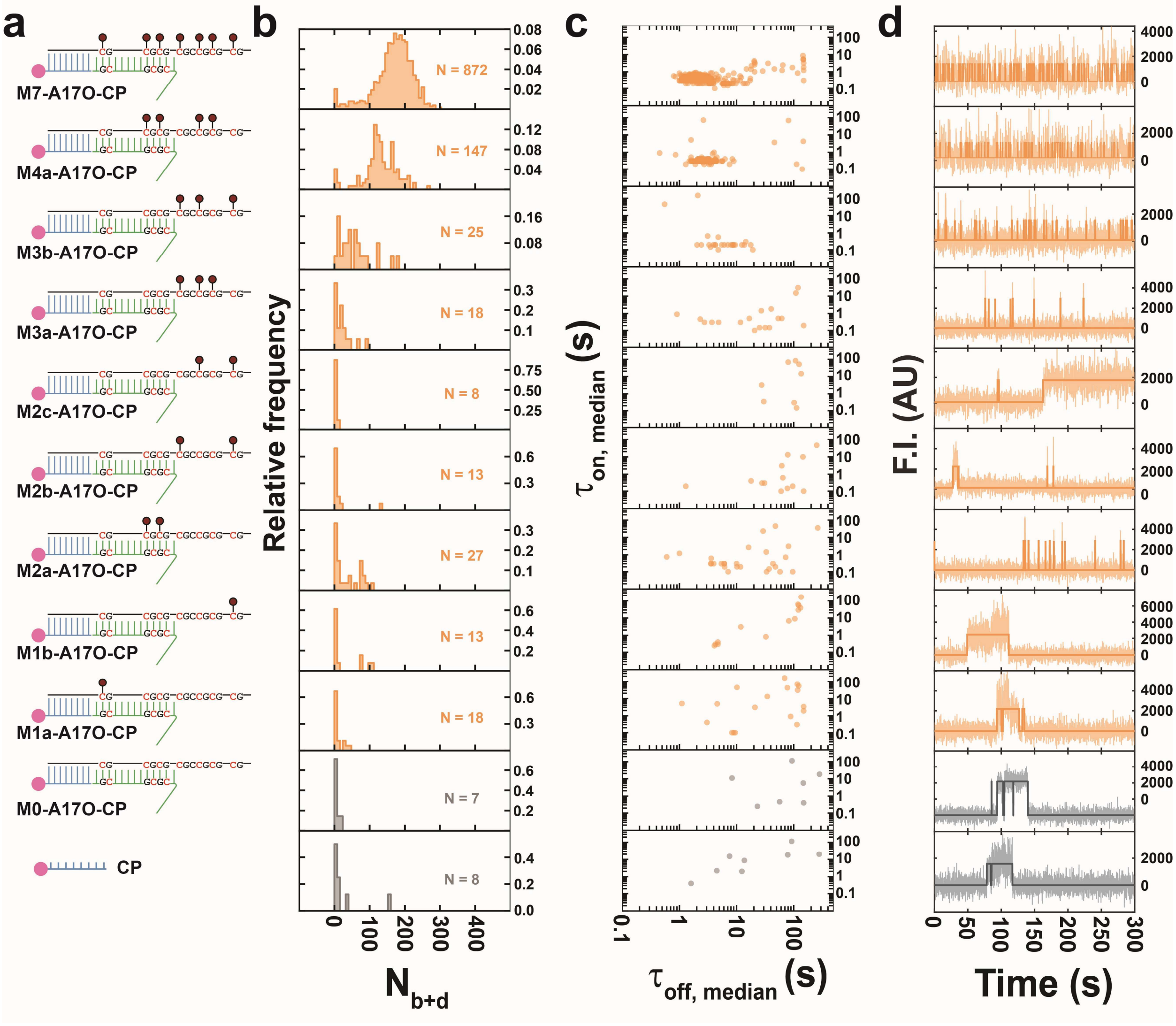


**Supplementary Fig. 12 | Binding kinetics of bifurcating hemimethylated DNA modulated by clustering of methyl-CpG sites. a** Constructs of M7-A17O-CP, M4a-A17O-CP, M3b--A17O-CP, M3a--A17O-CP, M2c--A17O-CP, M2b--A17O-CP, M2a--A17O-CP, M1b--A17O-CP and M1a-A17O-CP, M0-A17O-CP and CP. **b** $N_{b+d}$ distributions. N, the number of detected molecules of one field of view (FOV). **c** Median dwell time distributions. **d** Representative intensity-time traces. Semi-transparent lines in the background are raw traces and solid lines are idealized traces by hidden Markov modeling. F.I., fluorescence intensity; AU, arbitrary unit.

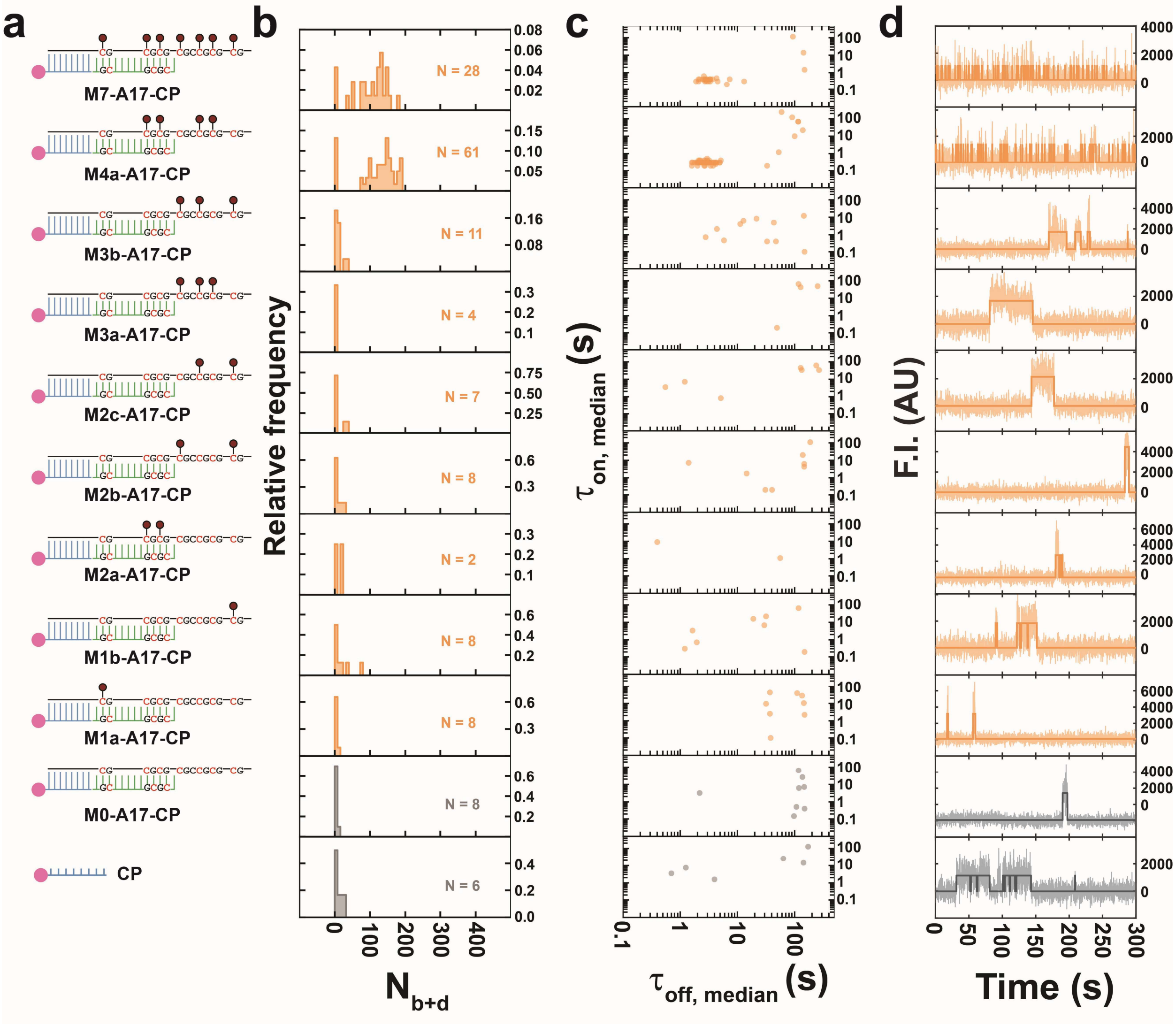


**Supplementary Fig. 13 | Binding kinetics of overhang-free partially exposed hemimethylated DNA modulated by clustering of methyl-CpG sites.** **a** Constructs of M7-A17-CP, M4a-A17-CP, M3b--A17-CP, M3a--A17-CP, M2c--A17-CP, M2b--A17-CP, M2a--A17-CP, M1b--A17-CP and M1a-A17-CP, M0-A17-CP and CP. **b** $N_{b+d}$ distributions. N, the number of detected molecules of one field of view (FOV). **c** Median dwell time distributions. **d** Representative intensity-time traces. Semi-transparent lines in the background are raw traces and solid lines are idealized traces by hidden Markov modeling. F.I., fluorescence intensity; AU, arbitrary unit.

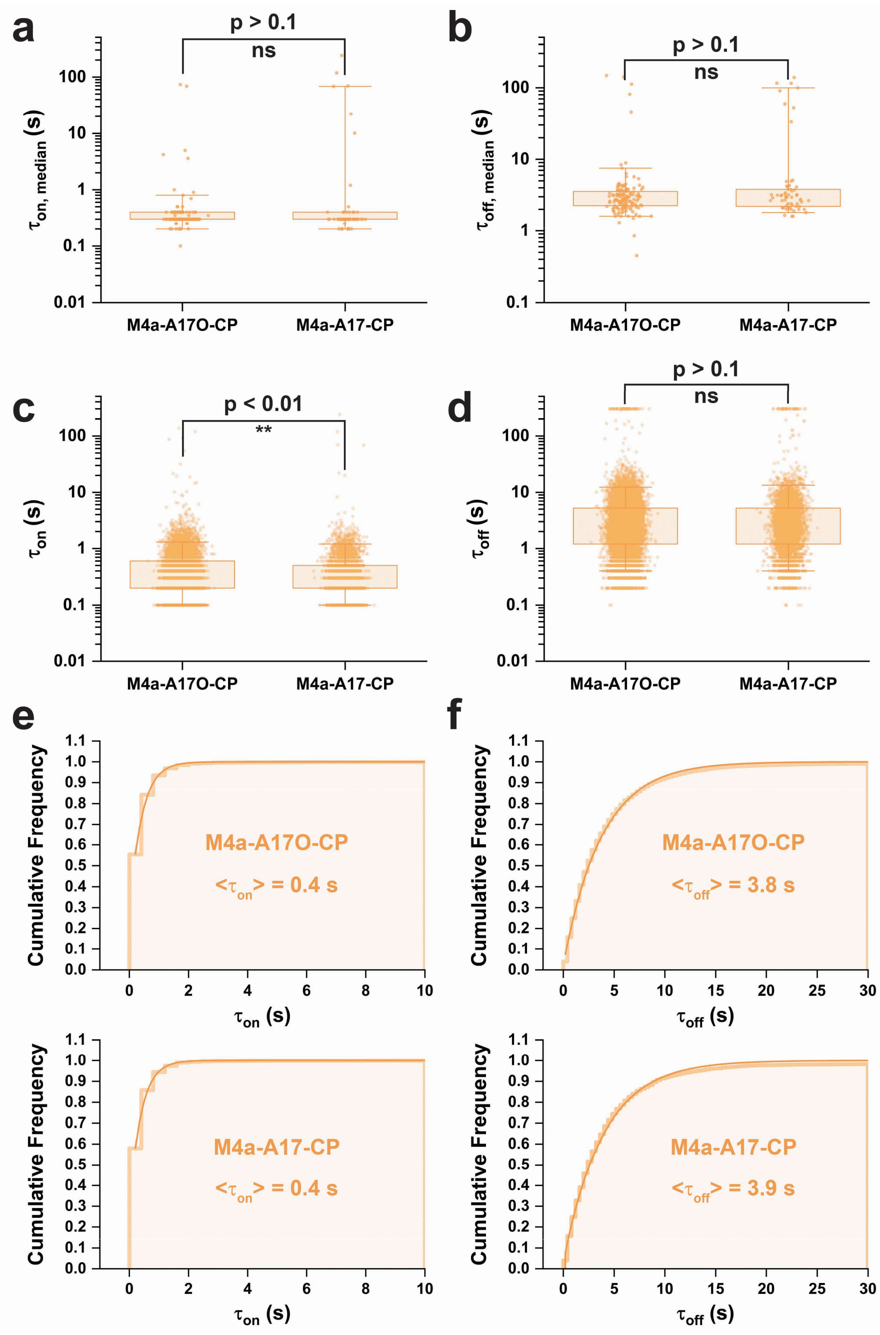


**Supplementary Fig. 14 | Dwell time comparison and mean dwell time calculations of M4a-A17O-CP and M4a-A17-CP. a,b** Boxplots of $\tau_{on,median}$ and $\tau_{off,median}$ distributions respectively. Boxes are drawn from Q1 to Q3 with whiskers from 5% percentile to 95% percentile. For hypothesis testing, all datasets are 96% winsorized to tolerate interference of outliers. P-values are assessed using two-tailed unpaired t-test. **c,d** Boxplots of $\tau_{on}$ and $\tau_{off}$ distributions respectively. Boxes are drawn from Q1 to Q3 with whiskers from 5% percentile to 95% percentile. P-values are assessed using Mann-Whitney U test. **e,f** Cumulative dwell time distributions and their exponential fittings of $\tau_{on}$ and $\tau_{off}$ of individual events of detected molecules. Step horizontal lines are cumulative frequency counts and solid curves are fitting curves. Mean dwell times, $\langle\tau_{on}\rangle$ and $\langle\tau_{off}\rangle$, are calculated by fitting a single-exponential decay function.

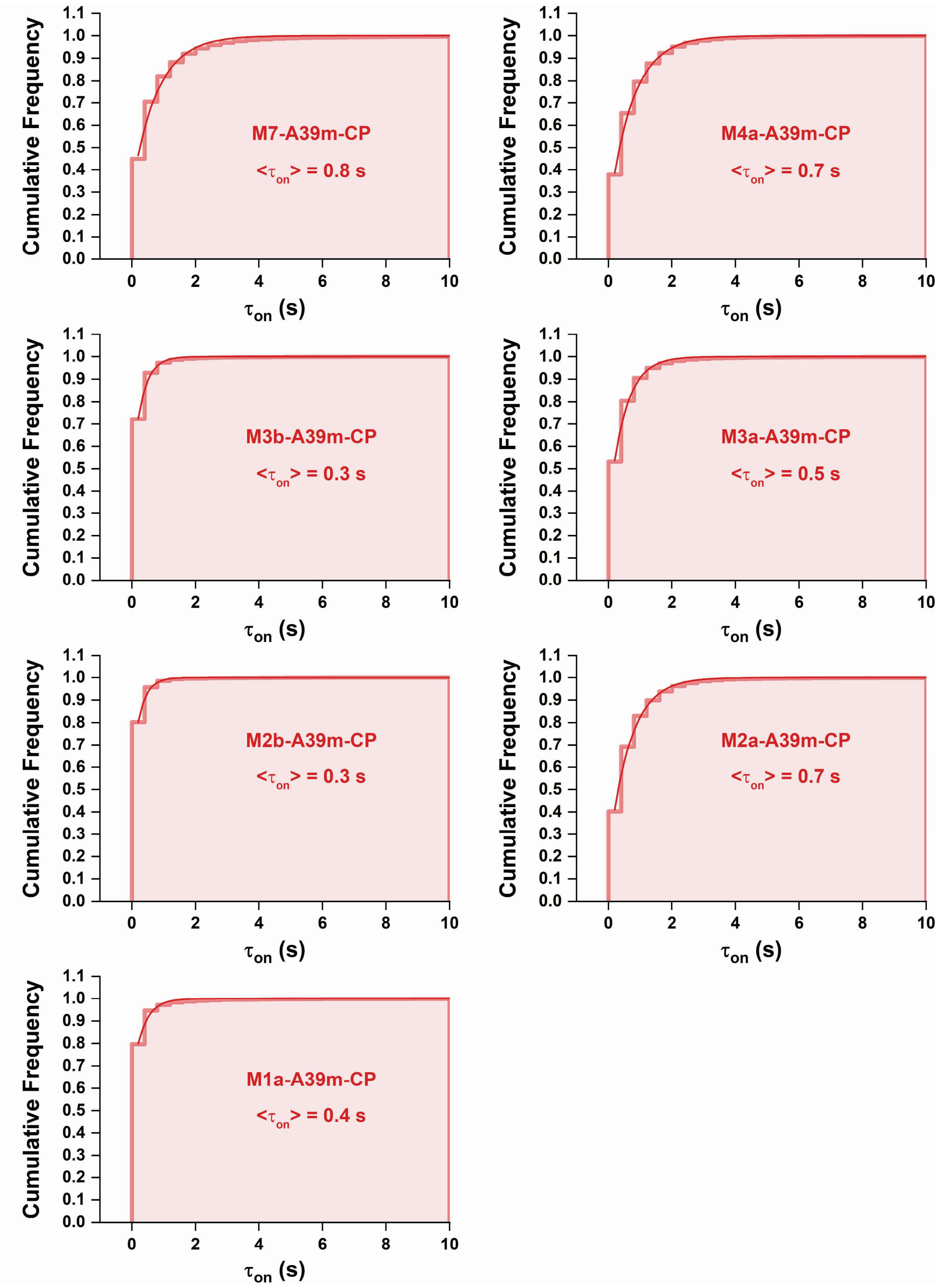


**Supplementary Fig. 15 | Mean bound time calculations of constructs in Fig. 4.** Cumulative dwell time distributions and their exponential fittings of $\tau_{on}$ of individual events of detected molecules. Step horizontal lines are cumulative frequency counts and solid curves are fitting curves. Mean bound time, $\langle\tau_{on}\rangle$, is calculated by fitting a single-exponential decay function.

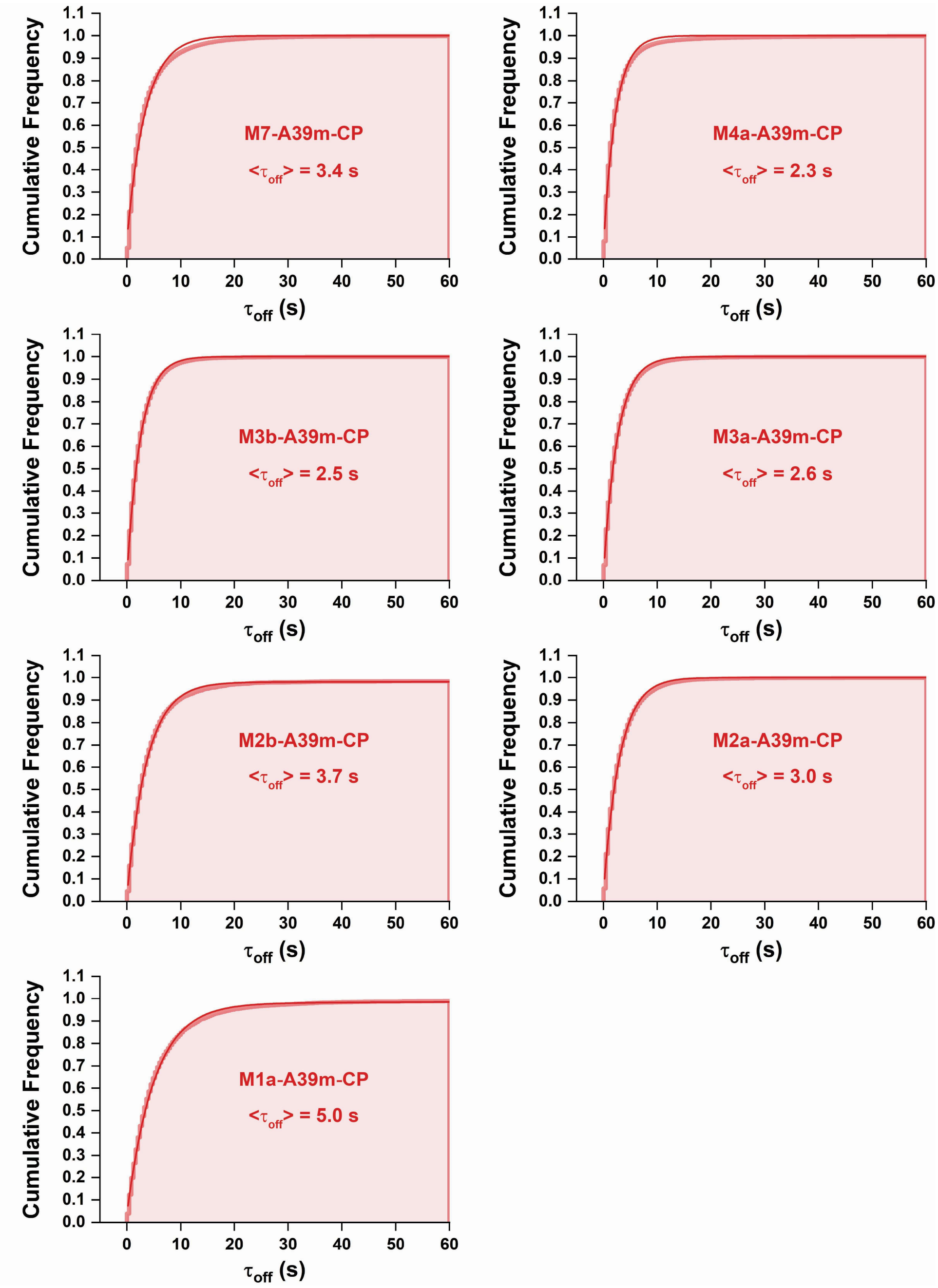


**Supplementary Fig. 16 | Mean unbound time calculations of constructs in Fig. 4.** Cumulative dwell time distributions and their exponential fittings of $\tau_{off}$ of individual events of detected molecules. Step horizontal lines are cumulative frequency counts and solid curves are fitting curves. Except for M2c-A39m-CP, M2b-A39m-CP and M1a-A39m-CP, mean unbound time, $\langle\tau_{off}\rangle$, is calculated by fitting a single-exponential decay function. For M2c-A39m-CP, M2b-A39m-CP and M1a-A39m-CP, mean unbound time, $\langle\tau_{off}\rangle$, is calculated by fitting a double-exponential decay function and taking only the primary component since the secondary component is marginal (< 5% compared to the primary component) and way too big (> 180 s).
